# The *Plasmodium falciparum* histone methyltransferase PfSET10 is dispensable for the regulation of antigenic variation and gene expression in blood stage parasites

**DOI:** 10.1101/2024.06.25.600636

**Authors:** Matthias Wyss, Abhishek Kanyal, Igor Niederwieser, Richard Bartfai, Till S. Voss

## Abstract

The malaria parasite *Plasmodium falciparum* employs antigenic variation of the virulence factor *P. falciparum* erythrocyte membrane protein 1 (PfEMP1) to escape adaptive immune responses during blood infection. Antigenic variation of PfEMP1 occurs through epigenetic switches in the mutually exclusive expression of individual members of the multi-copy *var* gene family. *var* genes are located in perinuclear clusters of transcriptionally inactive heterochromatin. Singular *var* gene activation is linked to locus repositioning into a dedicated zone at the nuclear periphery and deposition of histone 3 lysine 4 di-/trimethylation (H3K4me2/3) and H3K9 acetylation marks in the promoter region. While previous work identified the putative H3K4-specific methyltransferase PfSET10 as an essential enzyme and positive regulator of *var* gene expression, a recent study reported conflicting data. Here, we used iterative genome editing to engineer a conditional PfSET10 knockout line tailored to study the function of PfSET10 in *var* gene regulation. We demonstrate that PfSET10 is not required for mutually exclusive *var* gene expression and switching. We also show that PfSET10 is dispensable not only for asexual parasite proliferation but also for sexual conversion and gametocyte differentiation. Furthermore, comparative RNA-seq experiments revealed that PfSET10 plays no obvious role in regulating gene expression during asexual parasite development and gametocytogenesis. Interestingly, however, PfSET10 shows different subnuclear localization patterns in asexual and sexual stage parasites and female-specific expression in mature gametocytes. In summary, our work confirms in detail that PfSET10 is not involved in regulating *var* gene expression and not required for blood stage parasite viability, indicating PfSET10 may be important for life cycle progression in the mosquito vector or during liver stage development.

## Introduction

Malaria, a devastating infectious disease caused by protozoan parasites of the genus *Plasmodium*, is accountable for more than 200 million clinical cases and 600,000 deaths each year, mainly among young children and pregnant women in sub-Saharan Africa ^1^. Of the five *Plasmodium* species known to infect humans, *Plasmodium falciparum* is responsible for the vast majority of severe and fatal malaria outcomes. During blood infection in the human host, short-lived extracellular forms of the parasite (merozoites) invade red blood cells (RBCs), develop intracellularly through the so-called ring and trophozoite stages and then multiply via schizogony to release up to 32 daughter merozoites ready to invade new RBCs. These reiterative replication cycles are responsible for all malaria-related morbidity and mortality as well as for asymptomatic chronic infections in hundreds of millions of semi-immune individuals. Moreover, during each replication cycle a small proportion of parasites commit to sexual development and give rise to ring stage progeny that differentiate within twelve days and across five distinct morphological stages into mature female or male stage V gametocytes ^2^. Gametocytes are the only forms of the parasite capable of infecting the mosquito vector and are hence essential to secure onward transmission to other humans.

The high virulence of *P. falciparum* and its capacity to establish chronic infection are closely linked to the expression of *P. falciparum* erythrocyte membrane protein 1 (PfEMP1), a clonally variant antigen exposed on the surface of infected RBCs (iRBCs) ^3^. PfEMP1 is encoded by the *var* gene family and each parasite genome contains approx. 60 *var* gene paralogs, each encoding an antigenically distinct PfEMP1 protein ^4–7^. *var* genes are expressed in a mutually exclusive manner (singular gene choice), i.e. only a single *var* gene/PfEMP1 variant is expressed in individual parasites ^8^. Different PfEMP1 variants bind to different host receptors expressed on the surface of endothelial cells or uninfected RBCs ^3, 9^ As a result, iRBCs cytoadhere and sequester in the microvasculature of various organs, which on the one hand contributes strongly to severe malaria pathology and on the other hand spares iRBCs from being cleared in the spleen ^3^. Importantly, clonal antigenic variation of PfEMP1 allows parasites to also escape adaptive humoral immune responses and hence to sustain chronic blood infection and transmission for prolonged periods of time ^10–12^.

Antigenic variation is under epigenetic control and based on infrequent switches in mutually exclusive *var* gene activation that occur *in situ* in absence of any apparent recombination events ^8, 10–12^. All *var* genes are embedded in subtelomeric and some chromosome-internal heterochromatic domains that also contain numerous members of other clonally variant gene families (e.g. *rif*, *stevor*, *pfmc-2tm*, *phist*) ^13–16^. These transcriptionally silent domains are marked by histone 3 lysine 9 tri-methylation (H3K9me3) and heterochromatin protein 1 (HP1) ^14, 15, 17–20^. H3K9me3/HP1-enriched *var* genes are indeed transcriptionally silenced ^17–19^ and HP1 is essential for the maintenance and inheritance of the silenced state and mutually exclusive *var* gene transcription ^20^. In addition to HP1, the SIR2 and HDA2 histone deacetylases as well as the H3K36-specific histone lysine methyltransferase (HKMT) PfSET2 are also required for *var* gene silencing ^21–25^. In contrast to silenced *var* genes, the single active *var* locus lacks HP1/H3K9me3 and is marked instead by H3K9ac and H3K4me2/3 in the promoter region ^17–19^. The active *var* gene is transcribed by RNA polymerase II (RNA pol II) in ring stages and poised (repressed) in trophozoites and schizonts ^26^. Interestingly, the H3K4me2/3 and H3K9ac marks persist throughout the poised state suggesting they may serve as memory marks for re-activation of the same locus in the progeny ^17, 27^. Together, these studies firmly established that the silenced and active states of *var* genes are linked to distinct chromatin signatures that either prevent or permit transcription, respectively. However, the molecular mechanisms responsible for remodeling chromatin from a silenced into an active state exclusively at one single *var* gene locus remain largely obscure.

*P. falciparum* chromosome ends assemble into several clusters at the nuclear periphery, and chromosome-internal heterochromatin also localises to the perinuclear compartment ^14, 15, 23, 24, 28–30^. Results from fluorescence *in situ* hybridisation experiments (DNA-FISH/RNA-FISH) showed that *var* gene activation is linked to perinuclear locus repositioning ^14, 23, 24, 29–33^. Collectively, these studies suggested that mutually exclusive *var* gene activation takes place in a dedicated “*var* gene expression site” (VES) at the nuclear periphery, and corresponding models have been put forward by several laboratories ^14, 31–35^. Intriguingly, Volz and colleagues identified the HKMT PfSET10 as the first and, to our knowledge, so far only potential protein component of the proposed VES compartment ^35^. PfSET10 is conserved in apicomplexan parasites and harbors a predicted SET domain and a plant homeodomain (PHD) that displayed preferential binding to unmethylated and monomethylated H3K4 *in vitro* ^35^.

Immunoprecipitated PfSET10 catalysed the deposition of H3K4me2 and H3K4me3 marks *in vitro*, suggesting that PfSET10 is a H3K4-specific HKMT ^35^. Furthermore, PfSET10 localised exclusively to a distinct spot at the nuclear periphery ^35^. Notably, DNA-FISH combined with immunofluorescence assays (IFAs) on parasites selected or not for the mutually exclusive expression of particular *var* gene variants revealed that poised *var* genes localized in close spatial proximity to the PfSET10 focus, whereas this association was not observed when the corresponding genes where in a silenced state ^35^. Together, these findings established a model according to which PfSET10 specifically binds to the poised *var* gene promoter and deposits H3K4me2/3 marks to maintain and inherit a transcriptionally permissive chromatin state for re-activation of the same locus in the ring stage progeny ^35, 36^.

A recent study, however, challenged the proposed role of PfSET10 in the regulation of *var* gene expression ^37^. In contrast to Volz et al., who could not obtain *pfset10* knockout (KO) parasites ^35^, Ngwa and colleagues successfully generated a PfSET10 loss-of-function line expressing a C-terminally truncated PfSET10 mutant lacking the SET and PHD domains ^37^. These PfSET10 KO parasites showed no defects in cell cycle progression and replication, demonstrating that PfSET10 is in fact dispensable for asexual parasite proliferation ^37^. Importantly, quantitative reverse transcription PCR analyses of the entire *var* gene repertoire revealed no apparent differences in *var* gene expression patterns between PfSET10 KO and wild type control lines, suggesting that PfSET10 plays no major role in *var* gene regulation ^37^. However, since Ngwa et al. compared wild type parasites with straight *pfset10* KO parasites cultured independently for many generations, the authors could not rule out the possibility that any potential role for PfSET10 in parasite viability and/or mutually exclusive *var* gene transcription may have been compensated for by other H3K4-specific HKMTs ^37^. Furthermore, it remains unknown whether PfSET10 function is required for gametocytogenesis.

Here, we combined the rapamycin-inducible DiCre-dependent conditional knockout system ^38, 39^ with a drug selection-based approach to enrich for parasites expressing the same *var* gene ^40, 41^ in order to conduct a detailed functional analysis of PfSET10 in asexual parasites and gametocytes. In agreement with the work of Ngwa and colleagues, we show that PfSET10 is not essential for parasite proliferation and plays no apparent role in mutually exclusive *var* gene expression, switching, poising or epigenetic memory. Furthermore, we demonstrate that albeit PfSET10 shows a different localization pattern in early gametocytes compared to asexual parasites, and female-specific expression in late stage gametocytes, PfSET10 is not involved in regulating sexual commitment, gametocyte maturation or gamete activation. Lastly, comparative whole transcriptome analyses by RNA-seq show that PfSET10 plays no major role in regulating gene expression during asexual parasite proliferation and gametocyte differentiation.

## Results

### PfSET10 is not essential for asexual parasite replication

To facilitate the comprehensive functional analysis of PfSET10, we engineered a multipurpose DiCre-dependent inducible knockout cell line in the genetic background of 3D7/1G5DiCre ^39^. We first tagged *pfset10* at the 3’ end with a sequence encoding the promiscuous biotin ligase BirA* ^42^ fused to mNeonGreen (mNG) ^43^, calmodulin-binding-peptide (CBP) ^44^ and streptavidin-binding-protein (SBP) ^45^, with a synthetic intron containing a *loxP* element (*loxPint*) ^38^ inserted between the BirA* and mNG coding sequences (3D7/DiCre/SET10-BirA*xNCS) (Fig. S1). In a second step, we introduced another *loxPint* element into the *pfset10* coding sequence 1259 bp downstream of the start codon, yielding the cell line 3D7/DiCre/xSET10-BirA*xNCS (Fig. S1). As the functionalities of the BirA* and CBP/SBP affinity tag have not been employed in this study, we will refer to this cell line as 3D7/DiCre/SET10-mNG-iKO. Treatment with rapamycin (RAPA) induces DiCre-dependent recombination between the two *loxP* sites, resulting in excision of the majority of the *pfset10* coding sequence including the region encoding the SET and PHD domains (Fig. 1A and S1). Note that this recombination event also positions a premature stop codon downstream of the remaining *loxPint* sequence that prevents translation of the mNG tag (Fig. 1A and S1).With the 3D7/DiCre/SET10-mNG-iKO cell line established, we first investigated the subcellular localization of PfSET10-mNG in asexual blood stage parasites. Tightly synchronized parasites [0-4 hours post invasion (hpi)] were inspected by live fluorescence microscopy throughout the intra-erythrocytic developmental cycle (IDC). In agreement with previous observations ^35^, PfSET10-mNG localised exclusively to a unique spot at the nuclear periphery (Fig. 1B). Furthermore, PfSET10-mNG signals were first observed in late trophozoites at the 28-32 hpi time point (TP) (Fig. 1B) and then throughout schizogony until PfSET10-mNG signals disappeared in segmented schizonts beyond approx. 44 hpi (Fig. 1B).

**Figure 1.**
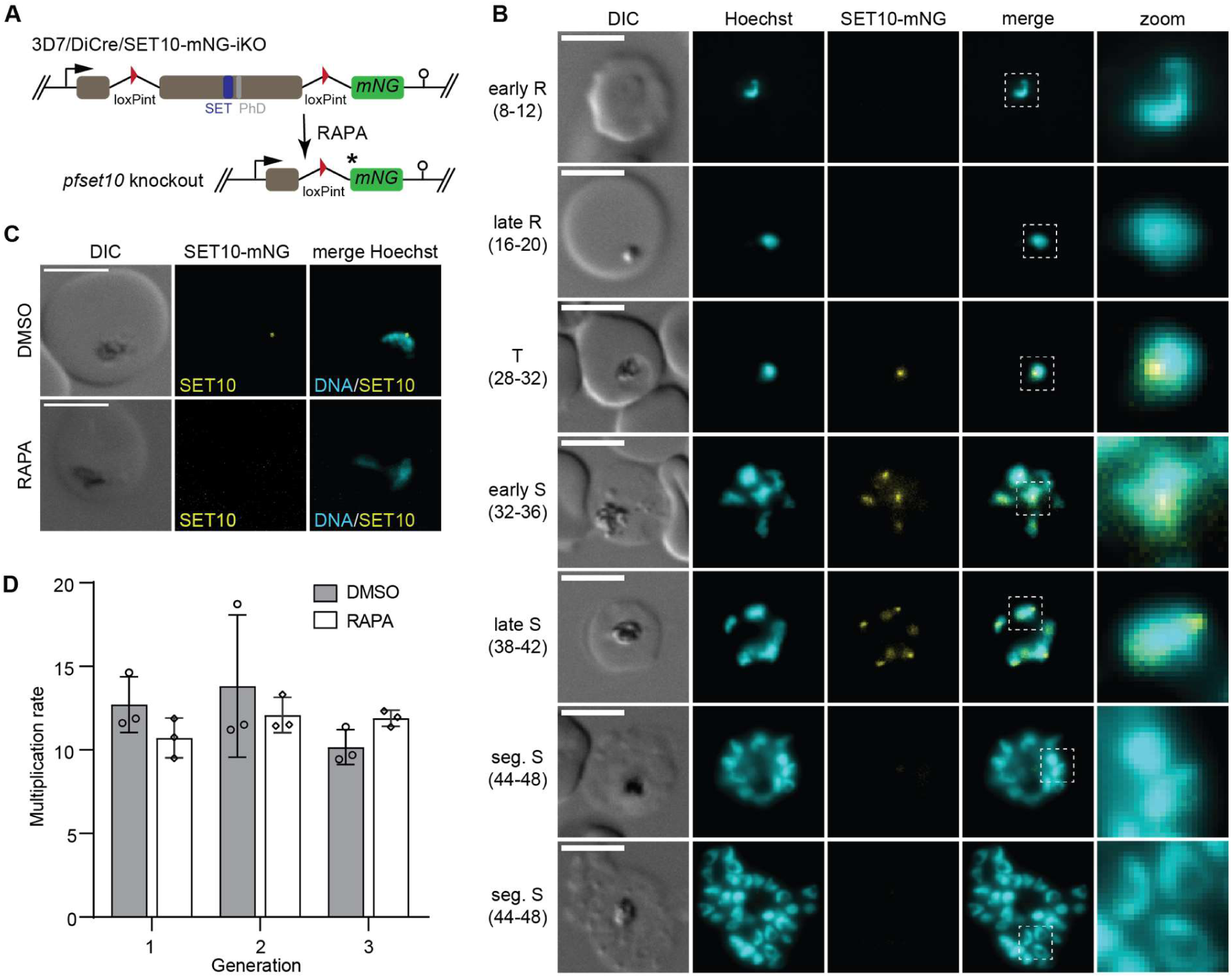
PfSET10 is not essential for asexual parasite replication. **(A)** Schematic of the 3D7/DiCre/SET10-mNG-iKO parasite line, before (top) and after (bottom) RAPA treatment. The relative position of the SET and PHD domain-encoding region is indicated. The asterisk denotes a STOP codon preventing expression of mNG after RAPA treatment. mNG, mNeonGreen. **(B)** Live cell fluorescence microscopy of PfSET10-mNG expression during asexual parasite development. Numbers in brackets indicate the age range (hpi, hours post invasion) of the synchronous parasite samples analysed at six TPs during the IDC. DNA was stained with Hoechst. DIC, differential interference contrast. R, ring; T, trophozoite; S, schizont, seg., segmented. Representative images are shown. Scale bar 5 µm. **(C)** Live cell fluorescence microscopy of PfSET10-mNG expression comparing control (DMSO) with PfSET10 KO (RAPA) parasites at 30-34 hpi. DNA was stained with Hoechst. DIC, differential interference contrast. Representative images are shown. Scale bar 5 µm. **(D)** Parasite multiplication rates measured by flow cytometry of SYBR Green I-stained control (DMSO) and PfSET10 KO (RAPA) parasites over three consecutive generations. Parasitemia was measured at 20-28 hpi. The means ±s.d. (error bars) of three biological replicates are shown, with circles representing individual values.

To confirm that RAPA treatment causes efficient DiCre-mediated deletion of *pfset10* and a the consequent loss of PfSET10-mNG expression, synchronized ring stage parasites (0-4 hpi) were split and treated with 100 nM RAPA or the DMSO solvent alone (control). As expected, PfSET10-mNG signals were undetectable in RAPA-treated parasites when inspected by live cell fluorescence microscopy (Fig. 1C). Furthermore, PCRs on genomic DNA (gDNA) isolated from schizonts confirmed the efficient excision of the *pfset10* gene (Fig. S1). Next, we performed parasite multiplication assays to test whether PfSET10 is essential for asexual blood stage parasite viability. Young ring stage parasites (0-8 hpi) were split and treated separately with RAPA and DMSO, respectively. Parasitemia was quantified by flow cytometry at 16-24 hpi over the course of three consecutive generations/invasion cycles. RAPA-treated parasites showed no signs of delayed growth or impaired multiplication compared to the control (Fig. 1D). These results are in agreement with recently published data obtained from mutant parasites expressing C-terminally truncated PfSET10 lacking the SET and PHD domains ^37^ and confirm that PfSET10 is not essential for asexual blood stage parasite proliferation *in vitro*.

### PfSET10 is not essential for *var* gene switching

It has been hypothesized that PfSET10 facilitates the maintenance and inheritance of a transcriptionally permissive chromatin state for re-activation of the active *var* locus in subsequent generations ^35^, but this proposed function has recently been challenged by Ngwa and colleagues ^37^. To investigate in more detail whether PfSET10 has any role in singular *var* gene choice and/or switching, we further modified the 3D7/DiCre/SET10-mNG-iKO cell line to be able to select for parasite populations expressing a particular *var* gene in a mutually exclusive manner. To achieve this, we made use of the concept underlying the selection-linked integration (SLI) approach ^46^. SLI is based on tagging genes at their 3’ end with a sequence encoding the 2A split peptide fused to a drug-selectable marker, which places expression of the marker under the control of the endogenous gene promoter and thus allows for the rapid selection of transgenic parasites via exposure to the corresponding selection drug ^46^. Recently, the SLI approach has successfully been used to select for parasites expressing particular members of clonally variant multigene families including *var* genes ^40, 41^. Here, we used CRISPR/Cas9-based gene editing to insert a sequence encoding a double TY tag (2xTY) fused to the 2A peptide followed by the blasticidin deaminase (BSD) drug-selectable marker at the 3’ end of *var* gene Pf3D7_0809100 (in the following called *v08*) in 3D7/DiCre/SET10-mNG-iKO parasites to obtain the cell line 3D7/DiCre/SET10-mNG-iKO/V08-2xTY^BSD^ (Fig. S2). To quantify the percentage of parasites expressing the V08-2xTY PfEMP1 variant prior to and post BSD-S-HCl selection, young ring stage parasites (0-8 hpi) were split and cultured separately in the constant absence (control) or presence of 2.5 ng/μl BSD-S-HCl. Exposure to BSD-S-HCl led to an initial decline in parasitemia and cultures resumed normal growth after two to three generations, suggesting successful selection of *v08*-expressing parasites. Immunofluorescence assays (IFAs) using α-TY1 antibodies were then performed on late trophozoites/early schizonts (28-36 hpi) and V08-2xTY-expressing cells were quantified based on the presence of punctate α-TY signals at the host RBC periphery consistent the localisation pattern of exported PfEMP1 (Fig. S3). In the BSD-selected 3D7/DiCre/SET10-mNG-iKO/V08-2xTY^BSD^ population, 90% of infected RBCs (iRBCs) were found to express V08-2xTY at the host cell membrane, while in the unselected population only 45% of parasites stained positive (Fig. 2A). This result shows that the 3D7/DiCre/SET10-mNG-iKO/V08-2xTY^BSD^ cell line can indeed be selected to express *v08-2xty* as the dominant *var* gene in most parasites in the population. The relatively high proportion of V08-2xTY-expressing parasites prior to BSD-S-HCl exposure was unexpected and may reflect preferential gene editing in parasites in which this locus was active and thus in a more accessible chromatin conformation at the time of transfection.

**Figure 2.**
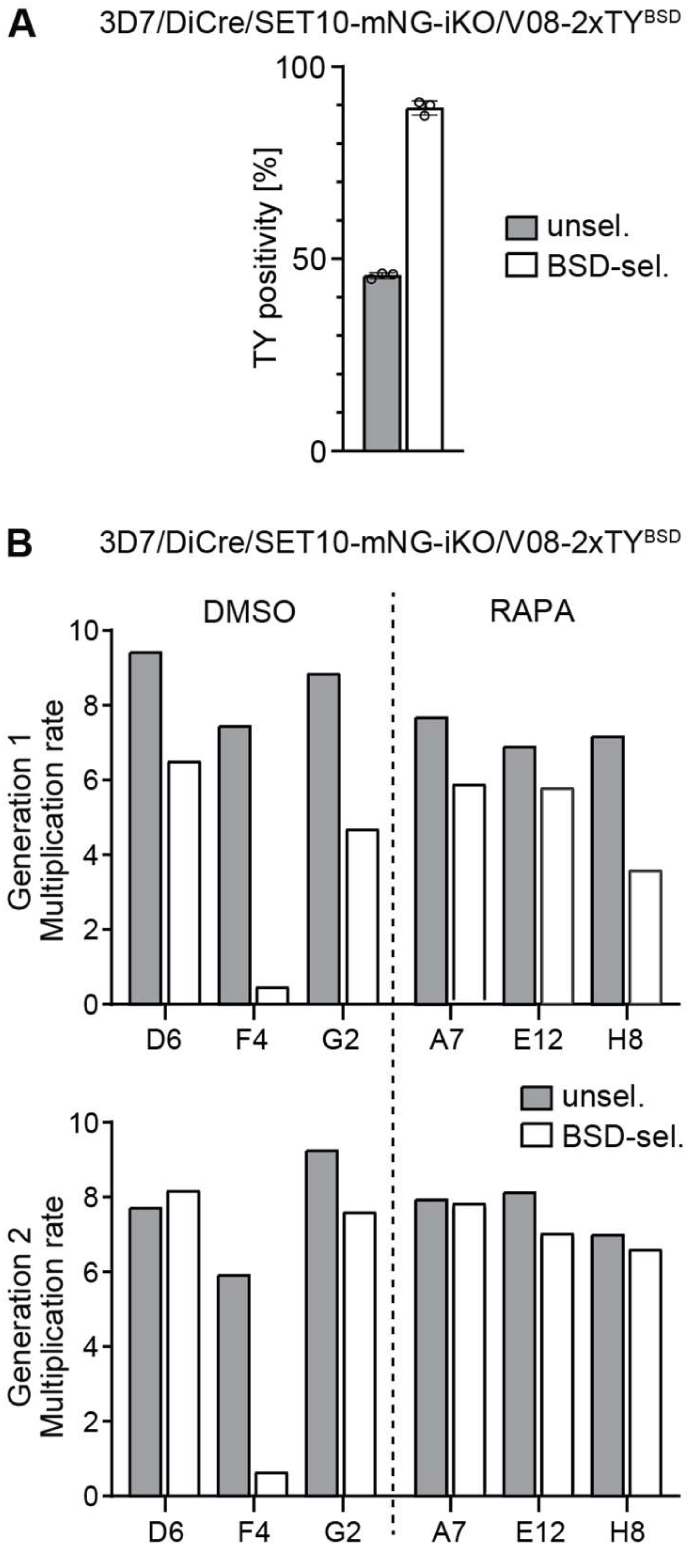
PfSET10 is not required for *var* gene switching. **(A)** Quantification of V08-2xTY-expressing cells in 3D7/DiCre/SET10-mNG-iKO/V08-2xTY^BSD^ parasites populations cultured in the absence (grey bar) or presence (white bar) of BSD-S-HCl pressure as assessed by IFAs using α-TY antibody at 32-40 hpi. The means ±s.d. (error bars) of three biological replicates are shown, with circles representing individual values. At least 150 individual iRBCs were counted for each condition and replicate. **(B)** Parasite multiplication rates measured by flow cytometry of three SYBR Green I-stained clonal lines each obtained from DMSO-treated (grey bars) and RAPA-treated PfSET10 KO parasite populations (white bars) after the first (top) and second (bottom) invasion cycles (generations 1 and 2). Parasitemia was measured at 20-28 hpi. Values represent the results from a single challenge experiment. Clone IDs are indicated on the x-axis.

Next, we investigated if PfSET10 is essential for *var* gene switching. To this end, 3D7/DiCre/SET10-mNG-iKO/V08-2xTY^BSD^ ring stage parasites (0-8 hpi) were split and one half was treated with RAPA to delete the *pfset10* gene. In the subsequent generation, parasites from both the DMSO control and RAPA-treated PfSET10 KO populations were subject to limiting dilution cloning. During clonal expansion, epigenetic memory ensures inheritance of *var* transcriptional states to the progeny, while infrequent switching events still occur. Hence, clonal populations are typically dominated by parasites expressing the same *var* gene that was active in the original clone and contain small subpopulations of parasites that express a different *var* gene ^47^. Here, we reasoned that if PfSET10 is essential for *var* gene switching, all parasites in clonal populations derived from RAPA-treated parsites would express the same *var* gene and would therefore be either entirely resistant or sensitive to BSD-S-HCl selection, depending on whether the original clone expressed *v08-2xty* or not, respectively. We therefore randomly selected three clonal populations each originating from the DMSO control and RAPA-treated PfSET10 KO cultures and exposed them to BSD-S-HCl pressure 20 generations (40 days) after seeding the clones. All six clonal populations showed reduced multiplication rates to various degrees in the presence of BSD-S-HCl compared to their paired control samples tested in the absence of drug (Fig. 2B). Furthermore, the replication rates of all clones, except the DMSO clone F4, recovered to the level of the corresponding control over the course of two generations (Fig. 2B). These results indicate that while five out of the six seeding clones likely expressed the *v08-2xty* gene (as expected from the high proportion of V08-2xTY-postive cells in the uncloned population prior to BSD-S-HCl selection), switching events took place during clonal expansion, showing that PfSET10 is not essential for functional *var* gene switching.

### PfSET10 is not required for epigenetic memory at the active *var* gene locus

The 3D7/DiCre/SET10-mNG-iKO/V08-2xTY^BSD^ line was also suitable to test the original hypothesis by Volz et al., who proposed that PfSET10 plays a crucial role in epigenetic memory by marking the active *var* gene for re-activation in the ring stage progeny ^35^. According to this model, parasites are expected to show increased *var* gene switching rates in absence of PfSET10 expression. To answer this question, we performed an RNA-seq time course experiment on paired DMSO control and RAPA-treated PfSET10 KO parasites preselected on BSD-S-HCl and then maintained in the absence of drug pressure for up to ten generations (Fig. 3A). In detail, 3D7/DiCre/SET10-mNG-iKO/V08-2xTY^BSD^ parasites were first selected on BSD-S-HCl to enrich for parasites expressing the *v08-2xty* gene. This bulk culture was then split into three subcultures. Culture 1 was kept under drug selection pressure (baseline reference), culture 2 was relieved from BSD-S-HCl pressure (DMSO control) and culture 3 was relieved from BSD-S-HCl pressure and treated with RAPA to delete the *pfset10* gene. After two additional generations, total RNA samples were collected from synchronous ring stages (10-16 hpi) in biological triplicates for the baseline reference (“R-WT-G0”), DMSO control (“R-WT-G2”) and RAPA-treated cultures (“R-KO-G2”). In addition, corresponding samples were collected again eight generations later for both the DMSO control (“R-WT-G10”) and RAPA-treated parasites (“R-KO-G10”) (Fig. 3A). As expected, *v08-2xty* was the only dominantly expressed *var* gene in the BSD-S-HCl-selected reference sample (R-WT-G0), with all other *var* transcripts showing low to very low normalized read counts, in agreement with the IFA-based quantification of V08-2xTY-positive iRBCs and consistent with mutually exclusive expression of *v08-2xty* in the vast majority of parasites (Figs. 3B and S4, Dataset S1). Upon removal of drug pressure, *v08-2xty* remained the dominant *var* transcript over the course of ten generations in both the DMSO control and RAPA-treated PfSET10 KO parasites. Some degree of *var* switching was evident over time in both populations as we observed increased transcript levels for several *var* genes ten generations post BSD-S-HCl removal compared to the BSD-S-HCl-selected reference sample (Figs. 3 and S4, Dataset S1). Together, these results demonstrate that PfSET10 is not required to maintain *var* genes in their active state across generations and confirms that PfSET10 plays no major role in regulating *var* gene switching.

**Figure 3.**
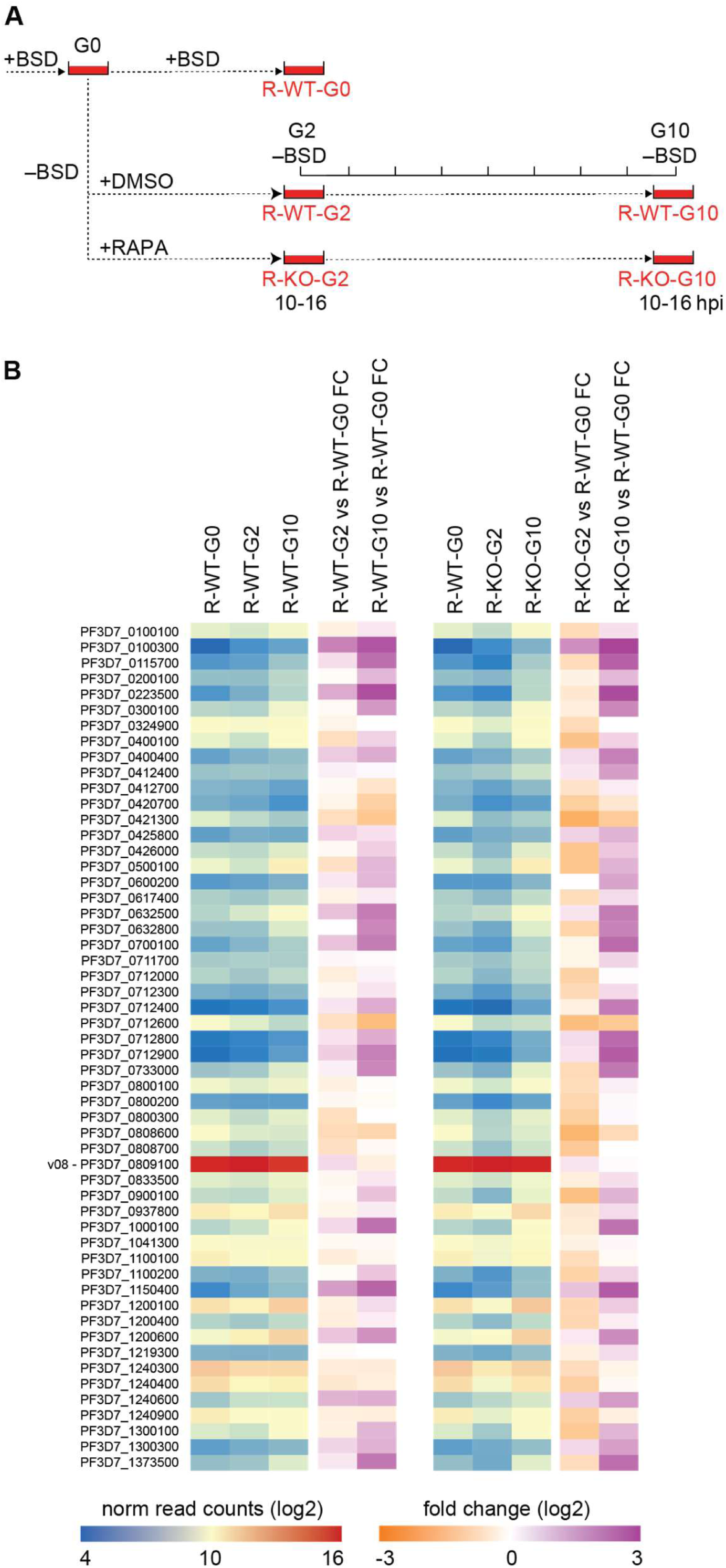
**PfSET10 is not required for mutually exclusive *var* gene expression and switching**. **(A)** Schematic explaining the sample collection workflow applied to quantify *var* gene transcripts by RNA-seq in DMSO control and RAPA-treated 3D7/DiCre/SET10-mNG-iKO/V08-2xTYBSD synchronous ring stage parasites two and ten generations after removal of BSD-S-HCl selection pressure. G, generation; R, ring stages; WT, DMSO-treated control parasites; KO, RAPA-treated PfSET10 KO parasites; hpi, hours post-invasion. **(B)** Heatmap showing normalized read counts (blue-red) and fold change in expression levels (orange-purple) of all *var* genes as quantified by RNA-seq of 3D7/DiCre/SET10-mNG-iKO/V08-2xTY^BSD^ ring stage parasites (10-16 hpi). R-WT-G0, control parasites selected on BSD-S-HCl for *v08-2xty* expression. R-WT-G2 and R-WT-G10, DMSO-treated control parasites harvested two and ten generations after release from BSD-S-HCl selection pressure, respectively. R-KO-G2 and R-KO-G10, RAPA-treated PfSET10 KO parasites harvested two and ten generations after release from BSD-S-HCl selection pressure, respectively. Values represent the mean of three biological replicate RNA-seq experiments performed for each condition and TP. The dominant *v08* gene PF3D7_1221000 has been highlighted.

### PfSET10 is not involved in regulating gene expression in asexual blood stage parasites

Despite the lack of any discernable PfSET10 loss-of-function phenotypes in asexual blood stage parasites, we still investigated whether the absence of PfSET10 affects gene expression during the IDC. Therefore, in addition to the ring stage samples R-WT-G2 and R-KO-G2 (10-16 hpi), we conducted RNA-seq analysis of three further TPs harvested in the same generation from trophozoites (22-28 hpi; T-WT-G2, T-KO-G2), early schizonts (30-36 hpi; ES-WT-G2, ES-KO-G2) and late schizonts (38-44 hpi; LS-WT-G2, LS-KO-G2) (Fig. 4A, Dataset S2). Principal component analysis (PCA) revealed tight clustering of the DMSO-treated control and RAPA-treated PfSET10 KO triplicate transcriptomes for each of the four IDC stages, suggesting deletion of *pfset10* has negligible effect on transcription in asexual blood stage parasites (Fig. 4B). Furthermore, successful disruption of the *pfset10* gene in RAPA-treated parasites was confirmed by the complete absence of RNA-seq reads mapping downstream of the inserted loxPint element (position +1259 bp relative to the ATG start codon), whereas reads mapping upstream were detected at levels comparable to those obtained from DMSO control samples (Fig. S5). We first compared the mean *v08-2xty* expression levels between the control and RAPA-treated PfSET10 parasites at all four IDC TPs, which showed that PfSET10 is not required for repression/poising of the active *var* gene in trophozoites and schizonts (Fig. S5). Next, we identified differentially expressed genes (DEGs) for each of the four TPs separately using DEseq2 ^48^. This analysis revealed 34, 4, 51 and 6 DEGs in ring stages, trophozoites, early schizonts and late schizonts, respectively (log2 FC >1; adjusted p-value <0.01) (Fig. S6, Dataset S2). Almost of all these genes were downregulated in the PfSET10 KO samples, with only a few upregulated genes detected at the early schizont TP. Closer inspection of the DEGs demonstrated that almost none of these genes showed consistent up- or downregulation in consecutive TPs, with *pfset10* being the only gene significantly downregulated at all four TPs (note that the FC values for *pfset10* represent the ratio of truncated versus full-length transcripts and hence do accurately reflect the degree of *pfset10* transcript depletion in the PfSET10 KO samples) (Fig. S6). Furthermore, the vast majority of DEGs were detected with low average read counts, had minimal expression at the corresponding TP and/or peak expression in the preceding TP, and displayed only moderate differential expression (only four genes including *pfset10* with greater than four-fold change) (Fig. S6 and Dataset S2). Based on these observations, and the low number of DEGs detected overall, we conclude that these changes in gene expression are likely not a direct result of the lack of PfSET10 expression but rather due to slight differences in parasite stage composition between the matching TPs and/or unreliable quantification of low abundance transcripts.

**Figure 4.**
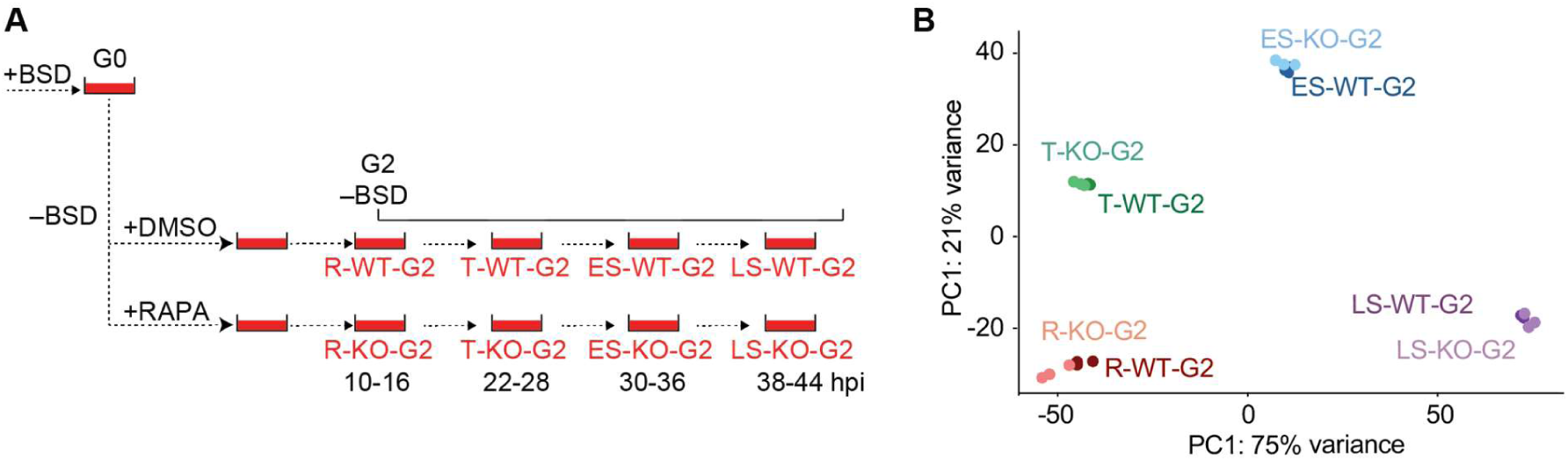
PfSET10 plays no major role in regulating gene expression in asexual blood dtage parasites. **(A)** Schematic explaining the sample collection workflow applied to quantify global gene expression by RNA-seq in synchronous DMSO control and RAPA-treated 3D7/DiCre/SET10-mNG-iKO/V08-2xTYBSD parasites at four time points during the IDC two generations after removal of BSD-S-HCl selection pressure. G, generation; R, ring stages; T, trophozoites; ES, early schizonts; LS, late schizonts; WT, DMSO-treated control parasites; KO, RAPA-treated PfSET10 KO parasites; hpi, hours post-invasion. **(B)** Principal Component Analysis plot of the triplicate transcriptomes determined by RNA-seq from synchronous 3D7/DiCre/SET10-mNG-iKO/V08-2xTY^BSD^ parasites at four TPs during the IDC, two generations after release from BSD-S-HCl selection pressure. R, ring stages (10-16 hpi); T, trophozoites (22-28 hpi); ES, early schizonts (30-36 hpi); LS, late schizonts (38-44 hpi). WT, DMSO-treated control parasites; KO, RAPA-treated PfSET10 KO parasites. G2, generation 2 after release from BSD-S-HCl selection pressure.

### PfSET10 is not required for gametocytogenesis but shows female-specific expression in late stage gametocytes

Since we did not observe any PfSET10 loss-of-function phenotypes in asexual blood stage parasites, we decided to also investigate PfSET10 expression and function during gametocytogenesis. However, as we observed poor gametocyte differentiation and lack of male gamete activation (exflagellation) with the 3D7/DiCre/SET10-mNG-iKO and 3D7/DiCre/SET10-mNG-iKO/V08-2xTYBSD lines, we re-engineered the PfSET10 inducible knockout cell line in NF54/DiCre parasites ^49^ using the same two-step gene editing approach explained above, yielding the NF54/DiCre/SET10-mNG-iKO line (Fig. S7). Successful DiCre-mediated excision of *pfset10* and lack of PfSET10-mNG expression in asexual

parasites was tested and confirmed by PCR on gDNA and live cell fluorescence microscopy, respectively (Fig. S7).

To monitor the expression and localization of PfSET10-mNG during gametocytogenesis, we induced sexual commitment in synchronous parasites using minimal fatty acid (mFA) medium as described ^50^. The ring stage progeny was maintained in serum-containing medium supplemented with 50 mM N-acetylglucosamine (GlcNAc) for six days to eliminate asexual parasites ^51^ and in absence of GlcNAc thereafter. Starting on day 2 of sexual development (stage I), gametocytes were inspected every second day until day 12 (stage V) using live cell fluorescence microscopy. Visualisation of the subpellicular microtubules with the live cell-compatible SPY555-tubulin stain was used to confirm stage-specific gametocyte morphology. We found that PfSET10-mNG was expressed from stage I gametocytes onwards (Fig. 5A). In contrast to asexual blood stage parasites, however, PfSET10-mNG was expressed throughout the Hoechst-stained area with several more densely stained spots observed (Fig. 5A). Interestingly, while PfSET10-mNG expression levels seemed to decrease in stage III gametocytes, from stage IV onwards only a subset of gametocytes expressed PfSET10-mNG (Fig. 5A). We hence suspected that PfSET10 might be expressed in a sex-specific manner in late stage gametocytes. To test this hypothesis, we performed dual-labeling IFAs on stage V gametocytes co-staining for PfSET10-mNG and Pfg377, a female-enriched osmiophilic body protein routinely used as a marker to differentiate female from male gametocytes ^52–55^. We indeed noticed that PfSET10-mNG-positive late stage gametocyte consistently also exhibited strong Pfg377 staining (Fig. 5B). To provide an unbiased quantification, we integrated the signal intensities originating from PfSET10-mNG and Pfg377 labeling using ImageJ as described previously ^56^. This analysis revealed that PfSET10-mNG was only expressed in gametocytes that also displayed strong Pfg377 signal intensities, indicating that in late stage gametocytes PfSET10 is indeed only expressed in females (Fig. 5C). Taken together, our results show that in gametocytes PfSET10 loses its exclusive association with the unknown perinuclear compartment observed in asexual parasites and localises much more broadly within the Hoechst-stained genetic material. In addition, PfSET10 assumes female-specific expression in late stage gametocytes.

**Figure 5.**
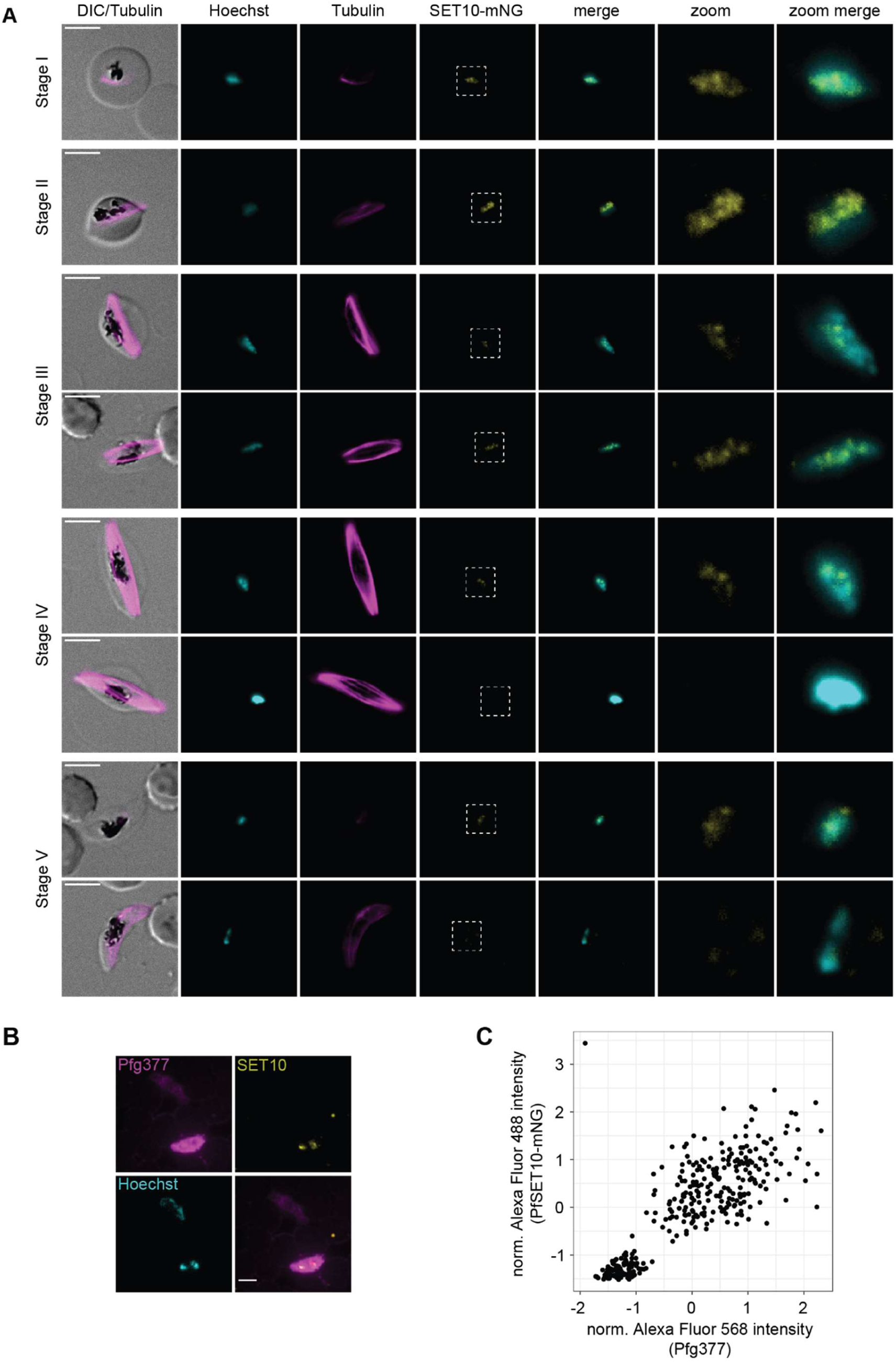
PfSET10 shows female-specific expression in late stage gametocytes. **(A)** Live cell fluorescence microscopy of NF54/DiCre/SET10-mNG-iKO gametocytes. PfSET10-mNG is visible from day 2 (stage I) onwards as multiple diffuse signals overlapping with the Hoechst-stained area. From day 8 (stage IV) onwards, a PfSET10-mNG negative subpopulation emerges. DNA was stained with Hoechst, subpellicular microtubules were stained with SPY555-tubulin. DIC, differential interference contrast. Representative images are shown. Scale bar 5 µm. **(B)** IFA overview image of NF54/DiCre/SET10-mNG-iKO stage V gametocytes (day 13) stained for the female-enriched marker Pfg377, PfSET10-mNG and Hoechst. Representative images are shown. Scale bar 5 µm. **(C)** Integrated Alexa Fluor 568 (Pfg377) and Alexa Fluor 488 (PfSET10-mNG) fluorescence signal intensity values resulting from automated image analysis of NF54/DiCre/SET10-mNG-iKO stage V gametocytes (day 13) stained with Hoechst, α-Pfg377/Alexa Fluor 568 and α-mNeonGreen/Alexa Fluor 488 antibodies. The scatter plot shows normalized integrated intensity values for Alexa Fluor 488 (PfSET10-mNG, y-axis) and Alexa Fluor 568 (Pfg377, x-axis) for 324 cells scored in a single experiment.

Next, we were interested in testing whether PfSET10 has any role to play in regulating sexual commitment, gametocyte development and/or gamete activation. We therefore induced excision of the *pfset10* gene PfSET10 in synchronized ring stage parasites (0-8 hpi) as explained above. RAPA-and DMSO-treated cultures were maintained for two additional generations before induction of sexual commitment using mFA medium ^50^. In the following generation, we performed IFAs on 30-38 hpi parasites using antibodies against the gametocyte-specific protein Pfs16 ^57^ to differentiate early stage I gametocytes from asexual trophozoites/early schizonts. We observed no difference in sexual conversion rates (i.e. the proportion of Pfs16-/DAPI-positive among all DAPI-positive iRBCs) between RAPA-treated PfSET10 KO and the DMSO-treated control populations (Fig. S8). Furthermore, gametocyte differentiation progressed normally without any noticeable delays in maturation or defects in morphology detected in PfSET10 KO gametocytes. Finally, to test whether the gamete activation capacity was impaired in absence of PfSET10, mature stage V gametocytes (day 13) were exposed to activation medium and female and male gamete activation was assessed by surface staining for Pfs25 (females) and exflagellation assays (males) using previously described assays ^58, 59^. We observed no defects in female or male gamete activation rates between PfSET10 KO and control gametocytes (Fig. S8).

Although gametocyte viability seemed unaffected in the absence of PfSET10, the broad distribution of PfSET10 in gametocyte nuclei and the female-specific expression in late stage gametocytes prompted us to test whether PfSET10 is involved in regulating gene expression during gametocytogenesis. Hence, we performed RNA-seq experiments on total RNA samples collected in biological triplicates from synchronous day 6 (stage III) and day 12 (stage V) PfSET10 KO and control gametocytes, obtained after mFA induction of sexual commitment and GlcNAc treatment as explained above (Dataset S3). In stage III gametocytes, the only differentially expressed gene identified was *pfset10* itself (Fig. S8). In stage V gametocytes, only six additional differentially expressed genes were detected including three clonally variant *rif* genes, all of which were 2-3-fold downregulated in PfSET10 KO gametocytes (Fig. S8). However, these few genes were expressed at low to very low levels in control gametocytes, so we consider their differential expression to fall within experimental noise. Hence, we conclude that PfSET10 has no major role in the regulation of gene expression in gametocytes.

In summary, despite the marked difference in PfSET10 nuclear localization patterns between asexual and sexual blood stage parasites and female-specific PfSET10 expression in late stage gametocytes, our comprehensive analysis did not identify any function for PfSET10 in regulating sexual conversion, gametocyte maturation or gamete activation.

## Discussion

With the study published by Volz and colleagues in 2012, PfSET10 has come into focus as a likely essential putative H3K4-specific HMKT and potential regulator of mutually exclusive *var* gene expression ^35^. Their data led to the compelling hypothesis that during schizogony, PfSET10 may be the enzyme depositing H3K4me2/3 marks specifically at the newly replicated copies of the poised *var* gene to tag these loci for re-activation in the subsequent ring stage progeny. Two recent studies, however, showed that disruption of the *pfset10* coding sequence had no obvious impact on parasite replication ^37, 60^. Furthermore, PfSET10 KO parasites displayed no marked difference in the overall *var* expression patterns and transcript levels compared to wild type control parasites ^37^. These results demonstrated that PfSET10 is indeed not essential in asexual blood stage parasites and questioned the proposed role for PfSET10 in regulating singular *var* gene choice.

Here, we revisited the expression dynamics, localization and function of PfSET10 in asexual and sexual blood stage parasites using RAPA-inducible DiCre-dependent PfSET10 knockout mutants. We employed this strategy to be able to identify PfSET10 loss-of-function phenotypes that may occur in direct response immediately after disruption of the *pfset10* locus and not obscured by potential compensatory mechanisms. Furthermore, to allow for a more precise assessment of any potential role PfSET10 may still play in the regulation of *var* gene expression, we additionally equipped the conditional PfSET10 mutant with a modified *var* gene locus (*v08-2xty*; Pf3D7_0809100) engineered to facilitate the BSD-S-HCl-based selection of parasites expressing *v08-2xty*/V08-2xTY in a mutually exclusive manner. Our results obtained from asexual parasites revealed that RAPA-treated PfSET10 KO parasites showed no reduction in multiplication rates over three subsequent generations. Clonal lines of control and RAPA-treated PfSET10 KO parasites could readily be selected for parasites expressing *v08-2xty*, showing that PfSET10 is not required for *var* gene switching. Furthermore, as shown by RNA-seq experiments, mutually exclusive transcription of *v08-2xty* was stably maintained in the absence of PfSET10 expression for ten generations with low levels of switching observed during this period, and this temporal *var* gene expression pattern was indistinguishable from that observed in the isogenic control population. Hence, stable inheritance of the active state of *var* genes is independent of PfSET10. Together, these findings are in agreement with those recently published by Ngwa et al. ^37^ and unequivocally demonstrate that PfSET10 is not essential for parasite replication and plays no role in regulating singular *var* gene choice and switching. Despite the now firmly corroborated lack of a functional link between PfSET10 and *var* gene regulation, the described close spatial association between the poised *var* gene locus and the perinuclear subcompartment to which PfSET10 localizes cannot be ignored ^35^. While from this point of view it is admittedly puzzling that a HKMT other than PfSET10 would seem to perpetuate the H3K4me2/3 epigenetic marks at the poised *var* locus throughout schizogony into the following generation, it is still conceivable that PfSET10 represents a coincidental component of the proposed perinuclear VES.

Our experiments performed during gametocytogenesis revealed that PfSET10 is fully dispensable for both female and male gametocyte differentiation and gamete activation. However, in contrast to the single distinct perinuclear signal observed in trophozoite and schizont nuclei, we found that PfSET10 localised broadly and to multiple more diffuse spots throughout the Hoechst-stained area in stage I-III gametocytes. This result indicates that either the unknown structure to which PfSET10 localizes to in asexual parasites dissociates at the onset of sexual differentiation or that PfSET10 simply loses its association with this compartment. Intriguingly, in stage IV and V gametocytes, PfSET10 signals disappeared in males and remained weakly detectable only in female gametocytes. The biological significance of this observation, if any, remains unknown. However, we can at least speculate that PfSET10 may deposit H3K4me3 modifications at certain loci specifically in the maternal genome to mark these genes for activation during the gamete-to-zygote/ookinete transition and/or subsequent oocyst and sporozoite development. Interestingly, a recent study indeed reported significantly elevated H3K4me3 enrichment levels in female gametocytes at genes that are upregulated in ookinetes and oocysts ^61^.

The highly restricted localization of PfSET10 in schizonts would point at a rather limited set of potential target genes. In gametocytes, the more widespread distribution of PfSET10 throughout the Hoechst-stained chromatin mass indicates that a larger part of the genome may be associated with PfSET10 in these stages. However, our comparative RNA-seq analysis across the IDC and stage III and V gametocytes did not uncover any marked changes in gene expression in PfSET10 KO compared to control parasites, which is consistent with the lack of any PfSET10 loss-of-function phenotypes observed in asexual or sexual blood stage parasites. Our finding that PfSET10 plays no apparent role in regulating gene expression in blood stage parasites may be explained in several ways. First, reduced H3K4me3 enrichment at potential PfSET10 target genes may not affect their expression levels. Previous studies have shown that H3K4me3 is generally associated with active genes but, unlike H3K9ac, H3K4me3 occupancy does not correlate with gene expression levels ^61, 62^. Second, the loss of PfSET10 expression may not affect the overall H3K4me3 landscape due to redundant activities of other HKMTs known or predicted to methylate H3K4 (PfSET1, PfSET4, PfSET6, PfSET7) ^63, 64^. Third, it is still unknown whether or not PfSET10 actually associates with chromatin and methylates H3K4 *in vivo*. Previous efforts aiming to identify PfSET10 target genes by chromatin immunoprecipitation followed by sequencing (ChIP-seq) failed ^35, 36^, and our own attempts were also unsuccessful. In *in vitro* histone/nucleosome methylation assays, a recombinant protein encompassing the SET and PHD domains was inactive, and while affinity-purified PfSET10 fractions obtained from parasite lysates did methylate H3K4, the authors could not exclude that this reaction may have been carried out by co-purifying enzymatic activities ^35^. Furthermore, the SET domain annotated in PfSET10 is not closely related to the conventional SET domain consensus signature ^35, 63^, which is why it has initially not been identified as a SET domain-containing protein in a Hidden Markov Model-based query of the parasite proteome ^63^ and was only later classified as a divergent apicomplexan-specific H3K4-specific HKMT ^35, 65^. Hence, in light of these circumstances, and in the absence of direct experimental proof, we cannot entirely disregard the possibility that PfSET10 may methylate targets other than H3K4 or may not even possess methyltransferase activity.

Despite our tailored reverse genetics approach and comprehensive set of experiments, we were unable to discover any function for PfSET10 in *P. falciparum* asexual parasite development, replication, gametocyte maturation, gamete activation and gene regulation under *in vitro* cell culture conditions. Furthermore, our data related to the investigation of *var* gene expression corroborate the recent findings made by Ngwa et al. ^37^ and demonstrate in greater detail that PfSET10 is not involved in the regulation of singular *var* gene choice, maintenance of epigenetic memory and *var* gene switching. Results obtained from *in vivo* screens of barcoded *P. berghei* mutant libraries followed throughout the entire life cycle demonstrated reduced growth rates for PbSET10 KO parasites during blood infection ^66^ and strongly impaired liver stage development ^67^. We conclude that PfSET10 will be similarly important for *P. falciparum* liver stage development and the gametocyte-producing NF54/DiCre/SET10-mNG-iKO line generated here provides a suitable tool to study PfSET10 function in this important phase of the parasite life cycle.

## Methods

### Plasmodium falciparum cell culture

1. *P. falciparum* parasites were cultured as described previously ^68^ using AB+ or B+ human red blood cells (Blood Donation Center, Zürich, Switzerland) at a hematocrit of 5%. The culture medium contained 10.44 g/l RPMI-1640, 25 mM HEPES, 370 μM hypoxanthine, 24 mM sodium bicarbonate, 0.5% AlbuMAX II (Gibco #11021-037), 0.1 g/l neomycin, and 2 mM choline chloride (Sigma #C7527). Parasites were synchronized using 5% sorbitol in ddH2O ^69^. To induce the DiCre-mediated recombination of *loxP* sites, parasites were treated for at least 4 hours with 100 nM RAPA ^39^. Cultures were gassed with 3% O2, 4% CO2 and 93% N2 and incubated at 37°C.

### Cloning of transfection constructs

CRISPR/Cas9 plasmids: Small guide RNAs (sgRNAs) were cloned into BsaI-digested CRISPR/Cas9 plasmids by ligation of annealed complementary oligonucleotides. pHF_gC_SET10-Cterm was created by ligating annealed sgRNA_SET10_Cterm_F/_R encoding the sgRNA target agacatgttttaaatagaca (790-809 bp downstream of the *pfset10* stop codon) into pHF_gC ^70^. pBF_gC_SET10-S420 was created by ligating annealed oligonucleotides sgRNA_SET10_S420_5_F/_R encoding the sgRNA target aatgcgaatcgaagaaaaag (1269-1289 bp downstream of the *pfset10* start codon) into pBF_gC ^70^. pYF_gC-V08-4 was created by ligating annealed sgRNA_v08-4_F/_R encoding the sgRNA target gatgaatgaattgttagaaa (52-33 bp upstream of the *v08* stop codon) into pYF_gC. Prior to this, the pYF_gC vector was generated by replacing the human dihydrofolate reductase (*hdhfr*) selection marker in pHF_gC with the yeast dihydroorotae dehydrogenase selection marker (*ydhodh*). For this purpose, we used Gibson assembly ^71^ to combine three PCR products into EcoRI/AgeI-digested pHF_gC: (1) the *P. falciparum* calmodulin promoter amplified from pHF_gC using primers pYF_1F/_R; (2) *ydhodh* amplified from pY_gC ^72^ using primers pYF_2F/_R; and (3) part of the yeast cytosine deaminase/uridyl phosphoribosyl transferase negative selection marker *yfcu* amplified from pHF_gC using primers pYF_3F/_R.

Synthetic sequence encoding the mNG-CBP-SBP (NCS) tag: Overlapping complementary template oligonucleotides were designed that covered the whole sequences to be synthesized. Corresponding sets of oligonucleotides were mixed and the assembled sequence amplified by PCR with outer primers. First, a sequence encoding a tandem affinity tag consisting of calmodulin-binding-peptide (CBP) ^44^ and streptavidin-binding-protein (SBP) ^45^ was generated using the template oligonucleotides CS1 to CS4 at a final concentration of 3 nM each and the PCR primers CS0_F and CS5_R at 300 nM each. Next, the PCR product was cloned by restriction ligation using BamHI/XhoI into pETA_MBP ^73^ to generate pCS. Second, a sequence encoding mNeonGreen (mNG) ^43^, codon-adjusted for *P. falciparum*, was synthesized using 18 template oligonucleotides (N1 to N18) at 30 nM each and the PCR primers N0_F and N19_R at 300 nM each. A Gibson assembly reaction using this PCR product together with PCR-amplified pCS (primers NCS_F/_R) was then used to generate pMBP_NCS containing the sequence encoding the fusion tag mNeonGreen-CBP-SBP (NCS).

Donor plasmids: Donor plasmids were generated by Gibson assembly of PCR products, and in one case in addition with overlapping oligonucleotides. One of the PCR products for each donor consisted of the pD backbone ^70^. pD_SET10BxNCS was designed to C-terminally tag PfSET10 with BirA* and the NCS tag. In addition, a loxPint element ^38^ was introduced into the *birA*\* sequence and a terminator was included prior to the 3’ homology box (HB), since the sgRNA target site 800 bp downstream of the *pfset10* stop codon will cause removal of the endogenous terminator upon homologues recombination. For this purpose, we assembled the pD backbone with seven PCR-products: (1) a 450 bp 5’ HB (spanning bps 6538-6987 at the 3’ end of the *pfset10* coding sequence) amplified from 3D7 gDNA using primers Blox_1F/_R; (2) part I of the *birA*\* sequence amplified from pARL2_REX2-mDHFR-FKBP-myc-2A-BirA*-L1-FRB-L2-GFP ^74^ using primers Blox_2F/_R; (3) loxPint amplified from pRex2:loxPint:gfp ^38^ using primers Blox_3F/_R; (4) part II of the *birA*\* sequence amplified from pARL2_REX2-mDHFR-FKBP-myc-2A-BirA*-L1-FRB-L2-GFP using primers Blox_4F/_R; (5) the 985 bp NCS tag sequence amplified from pMBP_NCS using Blox_5F/_R; (6) the *P. berghei dhfr* terminator amplified form pHF_gC (Filarsky et al., 2018) using primers Blox_6F/_R; and (7) a 673 bp 3’ HB (spanning bps 841-1142 downstream of the *pfset10* gene stop codon) amplified from 3D7 gDNA using primers Blox_7F/_R.

The donor plasmid pD_SET10xS420 contained a loxPint element within codon S420 of the *pfset10* coding sequence. In addition, the sequence stretch covering codons K413-G433 was recodonized. pD_SET10xS420 was constructed by assembly of the pD backbone with three PCR products: (1) a 527 bp 5’ HB (spanning bps 751-1236 of the *pfset10* gene) amplified from 3D7 gDNA using primers SET10_x_S420_5HB_F/_R; (2) loxPint amplified from pD_SET10BxNCS using primers SET10_x_S420_F/_R; and (3) a 496 bp 3’ HB (spanning bps 1300-1749 of the *pfset10* gene) amplified from 3D7 gDNA using primers SET10_x_S420_3HB_F/_R.

The donor plasmid pD_V08-2TY-2A-BSD contained a recodonized sequence encoding the C-terminus of the V08 PfEMP1 (Pf3D7_0809100) (L2097 to I2109). pD_V08-2TY-2A-BSD was constructed by assembly of the pD backbone with three PCR products and four overlapping complementary oligonucleotides: (1) a 604 bp 5’ HB (spanning bps 6657-7213 at the 3’ end of the *v08* gene) amplified from 3D7 gDNA using primers v08_1F/_R; (2) overlapping oligonucleotides introducing a sequence encoding the 2xTY tag fused to the T2A skip peptide (v08_2F/_R and v08_3F/_R; (3) the *bsd* selection marker sequence amplified from pBF-gC ^70^ using primers v08_4F/_R; and (4) a 719 bp 3’ HB (spanning the stop codon and bps 1-689 downstream of the *v08* gene) amplified from 3D7 gDNA using primers v08_5F/_R.

All oligonucleotides used for the cloning of transfection constructs are listed in Table S1.

### Transfection and selection of genetically modified parasites

*P. falciparum* parasite were co-transfected with 50 μg donor plasmid and 50 μg CRISPR/Cas9 transfection vector as previously described ^70^. The 3D7/DiCre/SET10-BirA*xNCS and NF54/DiCre/SET10-BirA*xNCS parasite lines were generated by co-transfection of pD_SET10BxNCS and pHF-gC_SET10-Cterm into 3D7/1G5DiCre ^39^ and NF54/DiCre ^49^, respectively. The 3D7/DiCre/SET10-mNG-iKO (3D7/DiCre/xSET10-BirA*xNCS) and NF54/DiCre/SET10-mNG-iKO (NF54/DiCre/xSET10-BirA*xNCS) parasite lines were generated by co-transfection of pD_SET10xS420 and pBF-gC_SET10-S420 into 3D7/DiCre/SET10-BirA*xNCS and NF54/DiCre/SET10-BirA*xNCS, respectively. The 3D7/DiCre/SET10-mNG-iKO/V08-2xTYBSD parasite line was generated by co-transfection of pD_v08-2TY-2A-BSD and pYF-gC_V08-4 into 3D7/DiCre/SET10-mNG-iKO.

To select for transgenic parasites, different selection drugs and exposure times were used depending on the drug resistance marker encoded on the CRISPR/Cas9 plasmids. For pHF-gC-derived vectors 5 nM WR99210 was added for six days, for pBF-gC-derived vectors 5 µg/ml BSD was added for ten days, and for pYF-gC-derived 1.5 μM DSM1 was added for ten days. Drug selection was first applied 24 hours after transfection. Selection drugs were removed after the exposure times indicated above and the cultures were maintained in the absence of drug pressure until stably propagating parasite populations were obtained. Successful gene editing was confirmed by PCR on gDNA. All oligonucleotides used for diagnostic integration PCR reactions are listed in Table S1.

### Limiting dilution cloning

Limiting dilution cloning was performed according to a previously described protocol (Thomas et al., 2016) with slight modifications. In brief, synchronous ring stage parasite cultures (0.01% hematocrit) were diluted to obtain 0.5 and 0.1 parasites per 200 µl. For both dilutions, 200 µl culture aliquots were transferred to separate 96-well cell culture plates (Corning #353072) and kept in a gassed airtight container at 37°C for 12 to 14 days without medium change. Subsequently, the plates were imaged using the Perfection V750 Pro scanner (Epson, Nagano, Japan) to identify wells containing a single plaque in the RBC layer. The content of positive wells was transferred individually to 5 ml cell culture dishes and cultured under normal conditions until a stably growing parasite culture was obtained.

### Preparation of parasite samples for live cell fluorescence microscopy

Parasite culture suspensions were incubated 5 µg/mL Hoechst (Merck #94403) for 20 min at 37°C protected from light to stain parasite DNA. Microtubule staining was performed with SPY555-tubulin (Lubio Science #SC203) for 1 hour at 37°C. The stained samples were washed once in phosphate-buffered saline (PBS) and mounted on microscopy slides using Vectashield (Vector Laboratories #H-1200-10).

### Immunofluorescence assays

Thin blood smears prepared from pelleted parasite cultures were fixed in ice-cold methanol or methanol/acetone (60:40) for 2 min and stored at -80°C until further use. The fixed cells were incubated for 1 hour in blocking solution (3% BSA in PBS) followed by 1 hour incubation with the following primary antibodies diluted in blocking solution: mouse mAb α-mNeonGreen (Chromotek #32f6-100), 1:500; mouse mAb α-Pfs16 ^75^, 1:500; rabbit α-Pfg377 ^52^, 1:1000; mouse mAb α-TY1 (Invitrogen #MA5-23513), 1:1000. After three washes with PBS, the following secondary antibodies diluted in blocking solution were applied for 45 min in the dark: Alexa Fluor 488-conjugated α-mouse IgG (Molecular Probes #A11001), 1:500; Alexa Fluor 568-conjugated α-rabbit IgG (Molecular Probes #A11011), 1:500. Cells were mounted using Vectashield antifade containing DAPI (Vector Laboratories #H-1200).

### Fluorescence microscopy

Microscopy was performed with the Leica DM5000 B fluorescence microscope (40x and 63x objectives) or the Leica Thunder 3D assay fluorescence microscope (63x objective) equipped with Leica K5 cMOS cameras. Both microscopes use the Leica application suite X software (LAS X version 3.7.5.24914). Image processing was performed with Fiji (ImageJ version 1.52n). Within each experiment, identical settings were used for both image acquisition and processing.

### Quantification of Pfg377 and SET10-mNG signals in late-stage gametocytes

To assess the sex-specific expression of PfSET10-mNG in stage V gametocytes (day 13), IFAs α-mNeonGreen and α-Pfg377 antibodies and DAPI were performed on methanol/acetone-fixed thin blood smears. Image acquisition with the 40x objective was followed by automated quantification of the fluorophore signal intensities emitted from the secondary antibodies (Alexa Fluor 488-conjugated α-mouse IgG, Alexa Fluor 568-conjugated α-rabbit IgG) and DAPI using a Fiji macro (ImageJ version 1.52n) as described previously ^56^. In brief, images were segmented based on the Pfg377 signal using the auto-threshold “Triangle”, which identified potential male and female gametocytes. Regions of interest (ROIs) were defined by size threshold (15-40 µm^2^), circularity parameter (0.1-0.75) and empirically determined integrated DAPI signal intensity threshold. ROIs located at the image edges were excluded and for the remaining ROIs (gametocytes), the integrated signal intensities in the GFP (PfSET10-mNG) and Texas Red (Pfg377) channels were measured. Intensity values were divided by the corresponding gametocyte size (segmentation area) and normalized using the “scale()” function ^76^ in R (version 4.3.2) and plotted against each other in a scatter plot.

### Quantification of parasite multiplication rates

Parasite multiplication rates were determined using flow cytometry. Synchronised parasite cultures (0-8 hpi, 0.2% parasitaemia) were split and treated for 4 hours with 100 nM RAPA or DMSO. Parasites were cultured with daily medium changes and under shaking conditions. After completion of the first invasion cycle, cultures were diluted to 0.1-0.3% parasitemia and cultured for one additional cycle. Parasitaemia was measured at the start of the experiment (day 1) and after one (day 3) and two (day 5) invasion cycles. Each sample was stained with 1x SYBR-Green I (Invitrogen #S7563), incubated for 20 min in the dark, and washed once in PBS. Using a MACS Quant Analyzer 10, 50,000 events per condition were measured and analyzed using the FlowJo_v10.6.1 software. The proportion of iRBCs was determined based on the SYBR Green signal intensity. The gating strategy is shown in Fig. S9.

### Induction of sexual commitment and gametocyte cultures

Sexual commitment was induced by exposing synchronized parasite cultures (1-1.5% parasitemia) at 16–24 hpi to minimal fatty acid medium (mFA) for 32 hours as described ^50^. The mFA medium contains 0.39% fatty acid-free BSA (Sigma #A6003) instead of 0.5% AlbuMAX II and 30 μM each of the two essential fatty acids oleic and palmitic acid (Sigma #O1008 and #P0500, respectively). After completion of the invasion cycle, the mFA medium was removed from the ring stage progeny (0-8 hpi) and replaced with culture medium containing 10% human serum (Blood Donation Centre, Zürich, Switzerland) instead of 0.5% Albumax II. Parasites were cultured for another 30 hours before methanol-fixed thin blood smears were prepared at 30–38 hpi and IFAs performed using mouse mAb α-Pfs16 and DAPI. Sexual conversion rates were determined by quantifying the proportion of Pfs16/DAPI-positive parasites among all DAPI-positive parasites.

For gametocyte cultures, parasites were induced for sexual commitment as described above and the mFA medium replaced with serum-containing medium supplemented with 50 mM N-acetylglucosamine (GlcNAc) for six consecutive days with daily medium changes to eliminate asexual parasites ^51^. Thereafter, gametocytes were cultured in the absence of GlcNAc. For daily medium changes, gametocyte cultures were placed on a heating plate (37 °C) to prevent gametocyte activation.

### Male and female gametocyte activation assays

Exflagellation assays were performed on mature stage V gametocytes (day 13) as previously described^58^. Briefly, gametocyte cultures were pelleted at 400 g for 3 min and the RBC pellet was resuspended in activation medium (serum-complemented culture medium supplemented with 100 µM xanthurenic acid) at room temperature. After 15 min, the number of exflagellation centres and RBCs per ml of culture were quantified using a Neubauer chamber and bright-field microscopy (40x objective). Gametocytemia was determined by visual inspection of Giemsa-stained thin blood smears prepared from the same cultures. The percentage of exflagellating gametocytes was calculated as follows: 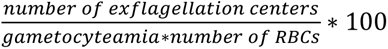.

Female gametocyte activation assays were performed on mature stage V gametocytes (day 13) as previously described ^59^. Briefly, gametocyte cultures were incubated overnight in activation medium containing mouse mAb α-Pfs25 primary antibodies (BEI Resources, #MRA-28) (1:2,000) and Alexa Fluor 594-conjugated α-mouse IgG secondary antibodies (Molecular Probes #A11005) (1:250) and then stained with SYBR Green (1:10,000). After washing with PBS, the samples (0.05% hematocrit) were transferred to imaging plates (Greiner # 655090; poly-D-Lysine, flat μClear bottom) and analyzed using the ImageXpress Micro wide-field high-content screening system (Molecular Devices) equipped with a Sola SE solid-state white light engine (Lumencor). Filter sets for SYBR Green [excitation (Ex): 472/30 nm, emission (Em): 520/35 nm] and Alexa Fluor 594 (Ex: 562/40 nm, Em: 624/40 nm) were used with exposure times of 40 and 50 ms, respectively. 121 sites per well were imaged using a Plan-Apochromat 40x objective (Molecular Devices, #1-6300-0412). Automated image analysis was performed using a modular image analysis workflow built with the MetaXpress software (version 6.5.4.532) to quantify rounded female gametes (i.e. the proportion of SYBR Green/Alexa Fluor 594 double-positive cells among all SYBR Green-positive cells) ^59^.

### Collection of total RNA for RNA-seq experiments

Synchronized parasites (0-6 hpi) cultured in the presence of 2.5 ng/μl BSD-S-HCl were split into three 10 ml cultures. Dish 1 (BSD control, baseline reference) was maintained under 2.5 ng/μl BSD-S-HCl selection. Dishes 2 (DMSO control, PfSET10 WT) and 3 (RAPA-treated, PfSET10 KO) were released from BSD-S-HCl pressure and treated with either the DMSO solvent alone or 100 nM RAPA, respectively. Dishes 1-3 were all cultured in parallel for two additional generations to allow for parasite expansion, and sorbitol synchronizations were performed to secure synchronous parasite populations for sample harvest. At the early ring stage (4-10 hpi), the DMSO- and RAPA-treated cultures were split into four equal 10 ml cultures each to be able to harvest four time points across the IDC (TP1, 10-16 hpi; TP2, 22-28 hpi; TP3, 30-36 hpi, TP4, 38-44 hpi), and one separate 10 ml dish each was maintained in culture for ten additional generations. The BSD control ring stage sample (R-WT-G0) was harvested alongside the DMSO- and RAPA-treated ring stage samples (R-WT-G2, R-KO-G2) (TP1), and trophozoite, early schizont and late schizont samples of the DMSO-treated (T-WT-G2, ES-WT-G2, LS-WT-G2) and RAPA-treated (T-KO-G2, ES-KO-G2, LS-KO-G2) cultures were harvested at the corresponding TPs 2-4 in the same IDC. For the DMSO- and RAPA-treated cultures, an additional 10-16 hpi ring stage sample each was harvested ten generations after release from BSD-S-HCl pressure (R-WT-G10, R-KO-G10). From each of these samples, total RNA was harvested from a 10 ml parasite culture (5% hematocrit; >3% parasitaemia) by resuspending the pelleted RBCs in 3 ml pre-warmed Trizol (Invitrogen #15596-018). After a 5 min incubation step at 37°C, the lysates were stored at -80°C until further use. After thawing, total RNA was purified using the Direct-zol RNA Miniprep Kit (ZYMO #R2050) including on-column DNA digest as per manufacturer’s instructions and RNA samples were eluted in 50 µl DNase/RNase-free water and stored at -80°C until further use.

For gametocyte samples, DMSO- and RAPA-treatment was performed on synchronous ring stage parasite cultures (0-8 hpi). After two additional invasion cycles, 20 ml parasite culture each was induced using mFA as explained above and the ring stage progeny was split into two 10 ml dishes, cultured in the presence of 50 mM GlcNAc for five days and total RNA samples were harvested on day 6 (D6-WT, D6-KO) and day 12 of gametocytogenesis (D12-WT, D12-KO), respectively.

### RNA-seq

Quality control (QC) of total RNA samples was assessed with the Agilent 4200 TapeStation system (Agilent Technologies) using High Sensitivity RNA ScreenTape analysis (Agilent, # 5067-5579) and all RNA samples were confirmed to be of high quality (mean RIN^e^ value =9.2 ±0.4 SD). RNA was quantified by fluorometry using the Qubit Flex Fluorometer (Thermo Fisher Scientific) and the Qubit RNA High Sensitivity Kit (Thermo Fisher Scientific, #Q32855). Sequencing libraries were prepared from 150 ng total RNA with the Illumina TruSeq Stranded mRNA library kit according to the manufacturer’s guidelines. QC of sequencing libraries was performed with the Agilent 5300 Fragment Analyzer system using the Standard Sensitivity NGS kit. Libraries were pooled and the pool cleaned up with 0.9 volumes SPRI beads (Beckman Coulter). The library pool was loaded in one Illumina NovaSeq 6000 S4 flow cell lane (concentration 220 pM) and sequenced to generate 101bp paired-end reads, with 50-80 million reads per sample generated.

### RNA-seq data analysis

The sequencing endline fastq files were quality checked using FASTQC (ver. 0.11.9). The reads were subsequently trimmed for quality (-q 30) and adapter removal using the Trim_Galore software (ver. 0.6.7). The trimmed reads were aligned onto the *P. falciparum* genome (ver. 64) available on www.plasmodb.org ^77^ using STAR aligner (ver. 2.7.10a). The third replicate of the DMSO control trophozoite samples was removed from the analysis owing to a low alignment rate. For the rest of the samples, the alignment BAM files were filtered for excluding reads mapping to mitochondrion- and apicoplast-derived RNAs and deduplicated using samtools package (ver. 1.15.1). The deduplicated reads were counted using the HTSeq package (ver. 0.11.1) [-count command with options: -s/reverse-stranded; -m/mode union]. The gene count tables for all samples were processed further for differential gene expression analysis using the DESeq2 package (ver. 1.36.0) in RStudio (ver. 4.2.2; ‘Innocent and Trusting’). Gene filters were set in DESeq2 to retain genes with more than 20 normalized counts in at least six samples. Pseudogenes and non-coding RNAs were removed from the list. Differential gene expression analysis for PfSET10 DMSO-treated control versus RAPA-treated PfSET10 KO samples was performed independently for asexual and sexual stage parasites and separately for each of the paired triplicate TP samples and genes with read counts ≥20. Significant differential expression was considered for genes with a mean log2 expression fold change of ≥1 (upregulated) or ≤-1 (downregulated) and Benjamini-Hochberg adjusted p-value <0.01. PCA plots were generated in DESeq2 with additional customization using ggplot2 package (ver. 3.4.4).

## Acknowledgements

Library preparation and sequencing were carried out in the Genomics Facility Basel of the University of Basel and the Department of Biosystems Science and Engineering, ETH Zurich, Switzerland. We are grateful to Sylwia Boltryk and Eilidh Carrington for technical assistance, and Pietro Alano and Robert Sauerwein for providing the α-Pfg377 and α-Pfs16 antibodies, respectively. The following reagent was obtained through BEI Resources, NIAID, NIH: Monoclonal anti-*Plasmodium falciparum* 25-kDa Gamete Surface Protein (Pfs25), Clone 4B7 (produced *in vitro*), MRA-28, contributed by Louis H. Miller and Allan Saul. This work was supported by a grant from the Swiss National Science Foundation (310031_184785).

## Author contributions

M.W., A.K., I.N., R.B. and T.S.V. conceived and designed the experiments. M.W. and I.N. performed the experiments. M.W., A.K. and T.S.V. analysed the data. T.S.V. and R.B. supervised the study. M.W. and T.S.V. wrote the manuscript. All authors contributed to the article and approved the submission.

## Data availability statement

All data generated or analysed during this study are included in this published article and its Supplementary Information files.

## Additional Information

The RNA-seq data published in this paper have been deposited on the GEO repository (GEO accession: GSE270631).

