## Supplementary Information for "The *Plasmodium falciparum* histone methyltransferase PfSET10 is dispensable for the regulation of antigenic variation and gene expression in blood stage parasites"

**This Supplementary Information file includes:**

- Figures S1-S9
- Table S1
- Legends for Datasets S1-S3

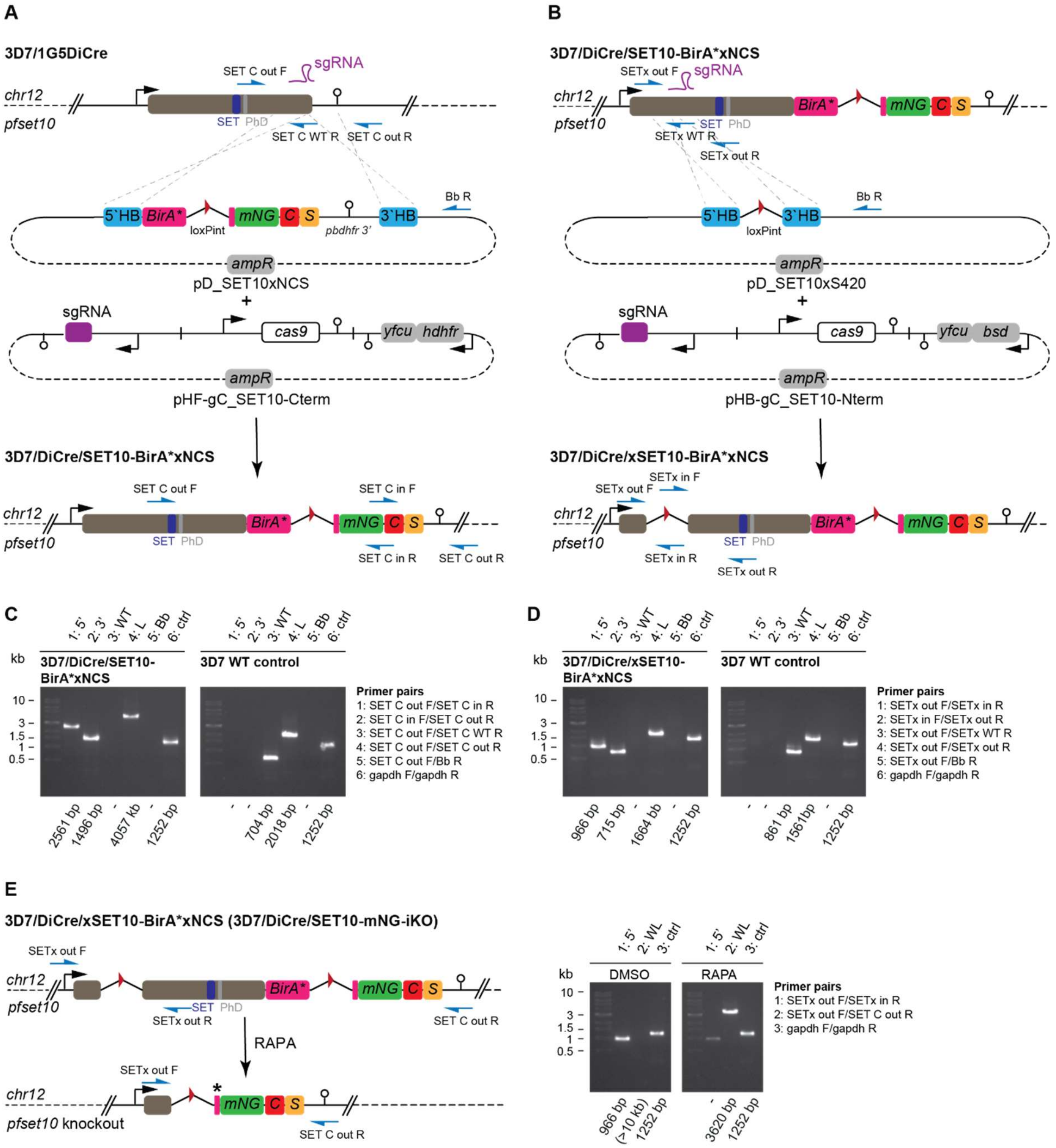


**Figure S1. Engineering and validation of the 3D7/DiCre/SET10-mNG-iKO line. (A)** Schematics of the *pfset10* wild type locus in 3D7/1G5DiCre parasites, the pD_SET10xNCS donor and pHF-gC_SET10-Cterm Cas9/sgRNA plasmids used to generate the 3D7/DiCre/SET10-BirA*xNCS line. The relative position of the SET and PHD domain-encoding region is indicated. Blue arrows indicate binding sites of primers used for diagnostic PCRs. HB, homology box; BirA*, promiscuous *E. coli* biotin ligase; mNG, mNeonGreen, C, CBP tag; S, SBP tag. loxPint, loxP element embedded in the *sera2* intron. **(B)** Schematics of the modified *pfset10* locus in 3D7/DiCre/SET10-BirA*xNCS parasites, the pD_SET10xS420 donor and pBF-gC_SET10-Nterm Cas9/sgRNA plasmids used to generate the 3D7/DiCre/xSET10-BirA*xNCS line. **(C-D)** Validation of the 3D7/DiCre/SET10-BirA*xNCS (C) and 3D7/DiCre/xSET10-BirA*xNCS (3D7/DiCre/SET10-mNG-iKO) (D) lines by diagnostic PCR on gDNA show correct genome editing and lack of parasites carrying the *pfset10* WT locus or plasmid integration. PCR primer pairs are indicated on the right, expected PCR fragment lengths at the bottom. 5'/3', PCRs detecting successful recombination of the 5' and 3' HBs, respectively; WT, PCR specific for the wild type *pfset10* sequence; L, PCR amplifying the entire locus from upstream to downstream of the 5' and 3' recombination sites; Bb, PCR detecting plasmid backbone integration; ctrl, control PCR amplifying the *gapdh* locus. **(E)** Schematic of the modified *pfset10* locus in 3D7/DiCre/xSET10-BirA*xNCS (here termed 3D7/DiCre/SET10-mNG-iKO) parasites before and after RAPA-induced DiCre-mediated excision of the floxed *pfset10* sequence (left). The asterisk denotes a STOP codon preventing expression of mNG after RAPA treatment. Diagnostic PCRs confirming successful excision of *pfset10* upon RAPA treatment (right).

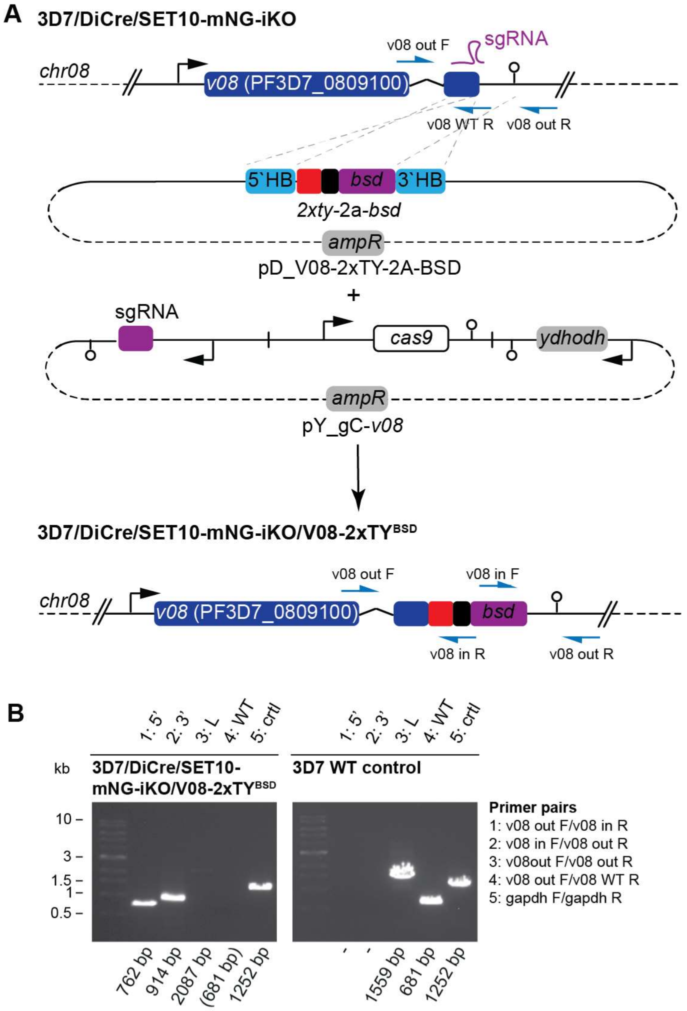


**Figure S2. 3D7/DiCre/SET10-iKO/v08-2xTY-2A-BSD cloning strategy and validation. (A)** Schematics of the wild type *v08* locus in 3D7/DiCre/SET10-mNG-iKO parasites, the pD_V08-2xTY-BSD donor and pY-gC_SET10-Nterm Cas9/sgRNA plasmids used to generate the 3D7/DiCre/SET10-mNG-iKO/V08-2xTY$^{BSD}$ parasite line. Blue arrows indicate binding sites of primers used for diagnostic PCRs. HB, homology box; *bsd*, blasticidin deaminase. Sequence elements encoding the 2xTY tag and 2A split peptide are represented by a red and black box, respectively. **(B)** Validation of the 3D7/DiCre/SET10-mNG-iKO/V08-2xTY$^{BSD}$ line by diagnostic PCR performed on gDNA shows correct genome editing and lack of parasites carrying the *v08* WT locus. PCR primer pairs are indicated on the right, expected PCR fragment lengths at the bottom. 5'/3', PCRs detecting successful recombination of the 5' and 3' HBs, respectively; L, PCR amplifying the entire locus from upstream to downstream of the 5' and 3' recombination sites; WT, PCR specific for the wild type *pfset10* sequence; ctrl, control PCR amplifying the *gapdh* locus. The missing fragment for the entire locus in 3D7/DiCre/SET10-mNG-iKO/V08-2xTY$^{BSD}$ parasites indicates plasmid integration.

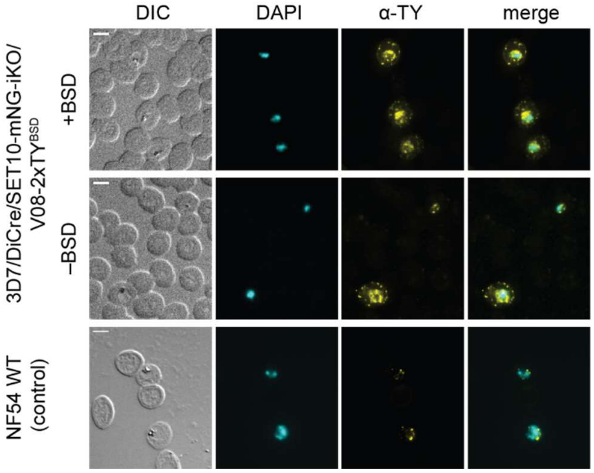


**Figure S3. BSD-S-HCl-based selection of 3D7/DiCre/SET10-mNG-iKO/V08-2xTY$^{BSD}$ parasites expressing the V08-2xTY PfEMP1 protein.** IFAs using α-TY antibodies to detect expression of V08-2xTY in RBCs infected with 3D7/DiCre/SET10-mNG-iKO/V08-2xTY$^{BSD}$ parasites at 32-40 hpi. The three iRBCs in the BSD-S-HCl-selected population show punctate V08-2xTY signals at the host cell periphery, consistent with exported PfEMP1 localisation (top panel). In the unselected population (middle panel), one iRBCs shows exported V08-2xTY signals whereas another cell does not express V08-2xTY but shows parasite-internal signals due to cross-reactivity of the α-TY1 antibody with an unknown parasite protein(s), which is consistently also observed in wild type NF54 parasites (bottom panel). Representative images are shown. DNA was stained with DAPI. DIC, differential interference contrast. Scale bar 5 µm.

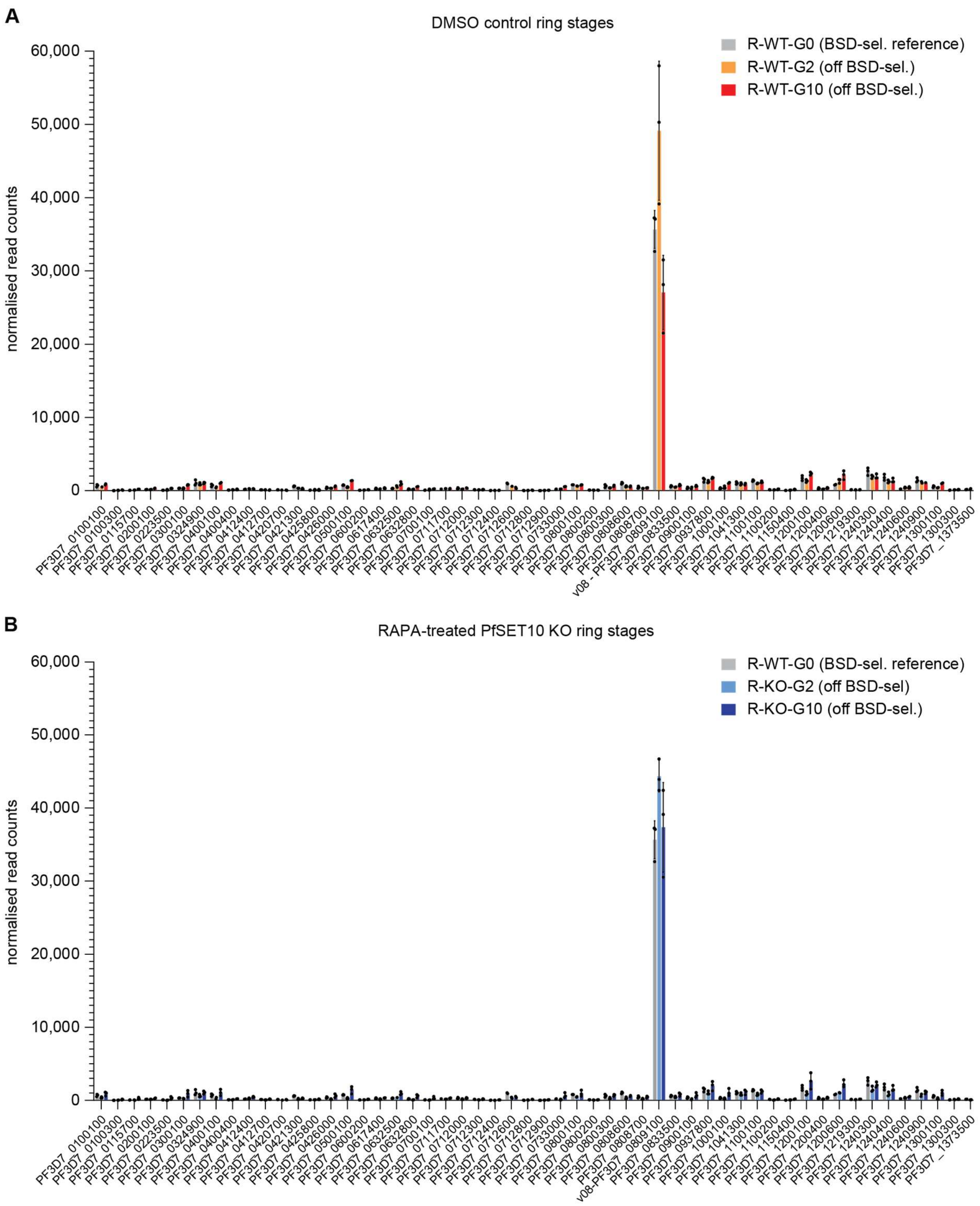


**Figure S4. *var* gene expression patterns followed over ten generations. (A, B)** Mean normalized read counts of all *var* genes as quantified by RNA-seq of 3D7/DiCre/SET10-mNG-iKO/V08-2xTY$^{BSD}$ ring stage parasites (10-16 hpi). R-WT-G0, control parasites selected on BSD-S-HCl for *v08-2xty* expression (A, B). R-WT-G2 and R-WT-G10, DMSO-treated control parasites harvested two and ten generations after release from BSD-S-HCl selection pressure, respectively (A). R-KO-G2 and R-KO-G10, RAPA-treated PfSET10 KO parasites harvested two and ten generations after release from BSD-S-HCl selection pressure, respectively (B). Values represent the mean of three biological replicate RNA-seq experiments performed for each condition and TP, with error bars representing the s.d. and circles representing individual values. The R-WT-G0 data in panels A and B are identical.

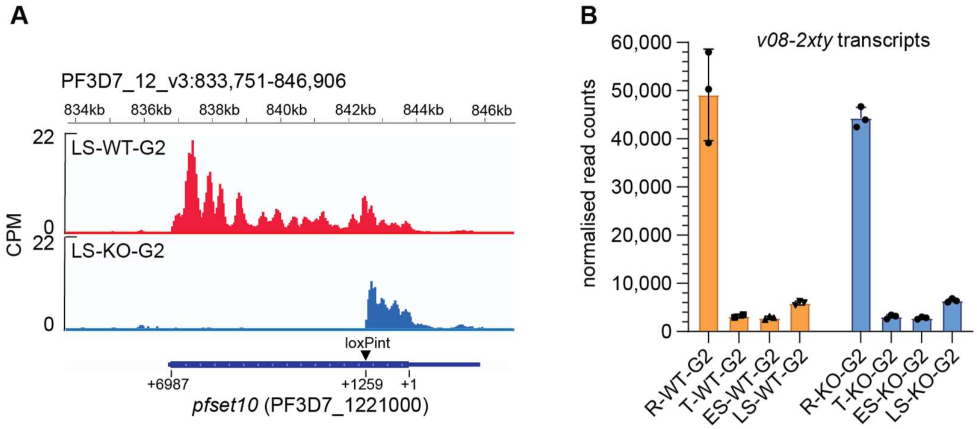


**Figure S5. (A)** Integrative Genomics Viewer browser tracks displaying coverage of reads mapped onto the *pfset10* locus in the DMSO-treated control (red, LS-WT-G2) and RAPA-treated PfSET10 KO (blue, LS-KO-G2) late schizont stage samples. In the PfSET10 KO sample, the *pfset10* reads are absent after the position of the inserted loxPint element (+1259 bp relative to the start codon). The y-axis represents counts per million (CPM) mapped reads across the locus in view with bin size of 50 bp. **(B)** Mean normalised *v08-2xty* read counts in control and PfSET10 KO parasites at four TPs during the IDC. R, ring stages (10-16 hpi); T, trophozoites (22-28 hpi); ES, early schizonts (30-36 hpi); LS, late schizonts (38-44 hpi). WT, DMSO-treated control parasites; KO, RAPA-treated PfSET10 KO parasites. G2, generation 2 after release from BSD-S-HCl selection pressure. Values represent the mean of three biological replicate experiments performed for each condition and TP, with error bars representing the s.d. and circles representing individual values.

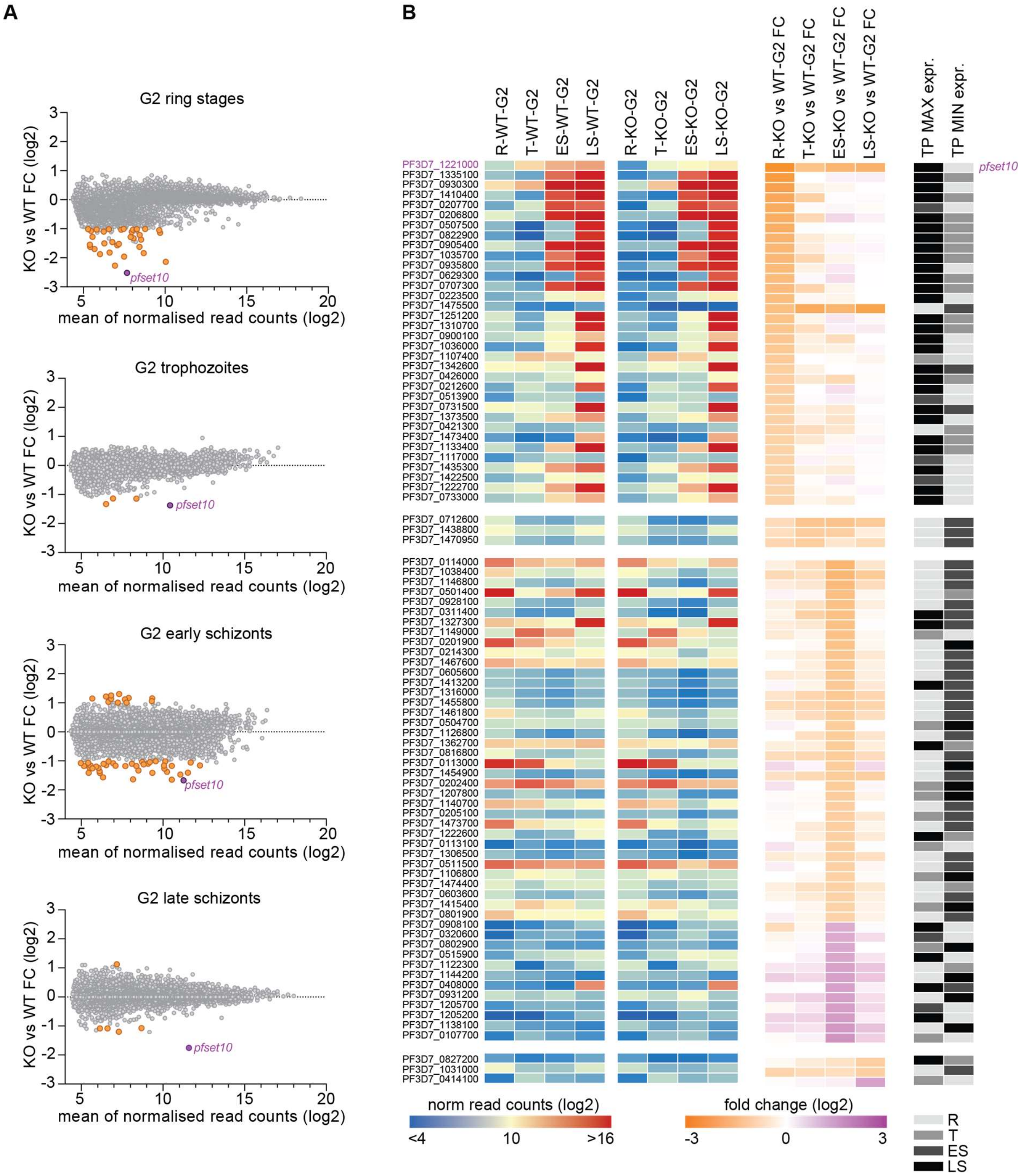


**Figure S6. Differential gene expression by DESeq2 analysis between 3D7/DiCre/SET10-mNG-iKO/V08-2xTY$^{BSD}$ DMSO control and RAPA-treated PfSET10 KO parasites at four TPs during the IDC.** (**A**) MA plots showing mean normalized read counts (x-axis) plotted against the fold change in gene expression between control and PfSET10 KO parasites (y-axis) for all genes and at four TPs during the IDC (ring stages, 10-16 hpi; trophozoites, 22-28 hpi; early schizonts, 30-36 hpi; late schizonts, 38-44 hpi). Coloured circles represent differentially expressed genes identified by DESeq2 analysis (log2 fold change (FC) ≥1; adjusted p-value <0.01). Values represent the mean of three biological replicate RNA-seq experiments performed for each condition and TP. WT, DMSO-treated control parasites; KO, RAPA-treated PfSET10 KO parasites. G2, generation 2 after release from

BSD-S-HCl selection pressure. (**B**) Heatmap showing the mean normalized read counts (blue-red) and fold change in expression (orange-purple) of all parasite genes identified as differentially expressed between control and PfSET10 KO parasites in at least one of the four IDC TPs based on DESeq2 analysis (log2 fold change (FC) ≥1; adjusted p-value <0.01). The heatmap on the right (grey-black) indicates the TPs of maximal and minimal expression levels during the IDC for each of the differentially expressed genes. The heatmap is partitioned into four subgroups, each containing the genes differentially expressed in ring stages, trophozoites, early schizonts and late schizonts, respectively, and sorted according to the fold change in gene expression in descending order. R, ring stages (10-16 hpi); T, trophozoites (22-28 hpi); ES, early schizonts (30-36 hpi); LS, late schizonts (38-44 hpi). WT, DMSO-treated control parasites; KO, RAPA-treated PfSET10 KO parasites. G2, generation 2 after release from BSD-S-HCl selection pressure. Values represent the mean of three biological replicate RNA-seq experiments performed for each condition and TP.

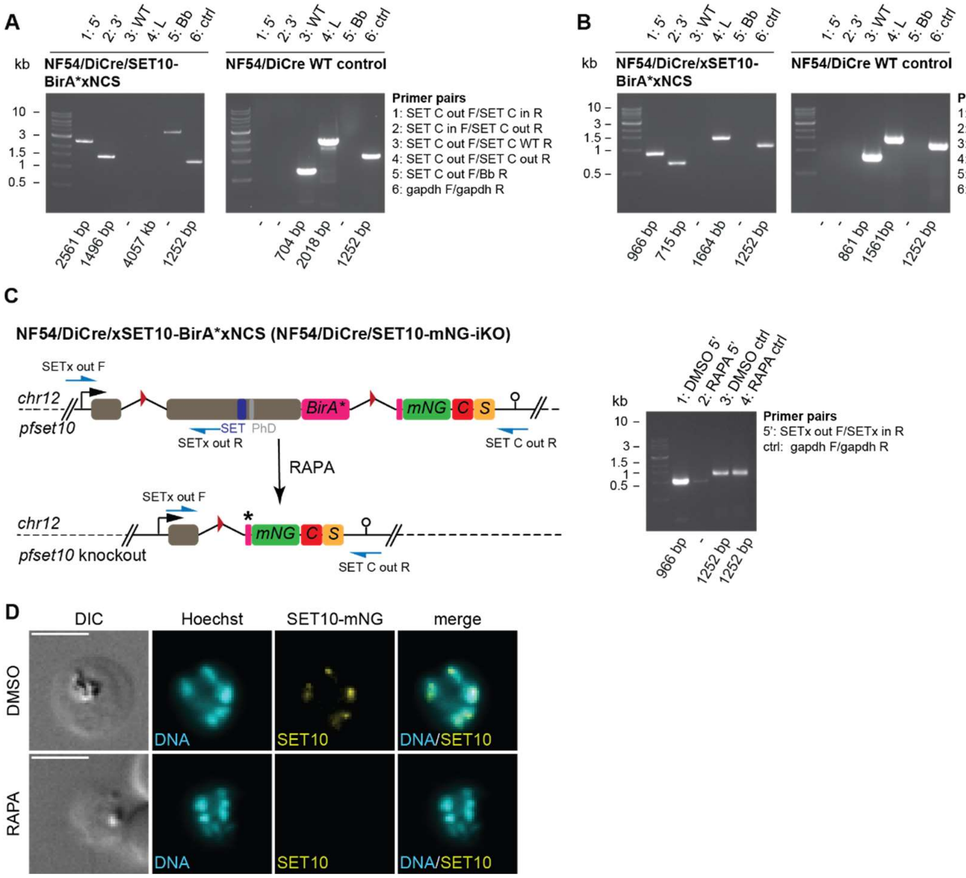


**Figure S7. Engineering and validation of the NF54/DiCre/SET10-mNG-iKO line. (A, B)** Validation of the NF54/DiCre/SET10xNCS (A) and NF54/DiCre/SET10-mNG-iKO (B) lines by diagnostic PCR on gDNA show correct genome editing and lack of parasites carrying the *pfset10* WT locus or plasmid integration. PCR primer pairs are indicated on the right, expected PCR fragment lengths at the bottom. 5'/3', PCRs detecting successful recombination of the 5' and 3' HBs, respectively; WT, PCR specific for the wild type *pfset10* sequence; L, PCR amplifying the entire locus from upstream to downstream of the 5' and 3' recombination sites; Bb, PCR detecting plasmid backbone integration; ctrl, control PCR amplifying the *gapdh* locus. Primer binding sites are shown in Fig. S1. **(C)** Schematic of the modified *pfset10* locus in NF54/DiCre/xSET10-BirA*xNCS (here termed NF54/DiCre/SET10-mNG-iKO) parasites before and after RAPA-induced DiCre-mediated excision of the floxed *pfset10* sequence (left). The asterisk denotes a STOP codon preventing expression of mNG after RAPA treatment. Diagnostic PCRs confirming successful excision of *pfset10* upon RAPA treatment (right). Primer binding sites are shown in Fig. S1. **(D)** Live cell fluorescence microscopy of PfSET10-mNG expression in schizonts (36-42 hpi) confirm efficient excision of *pfset10* in RAPA-treated NF54/DiCre/SET10-mNG-iKO parasites. DNA was stained with Hoechst. DIC, differential interference contrast. Representative images are shown. Scale bar 5 µm.

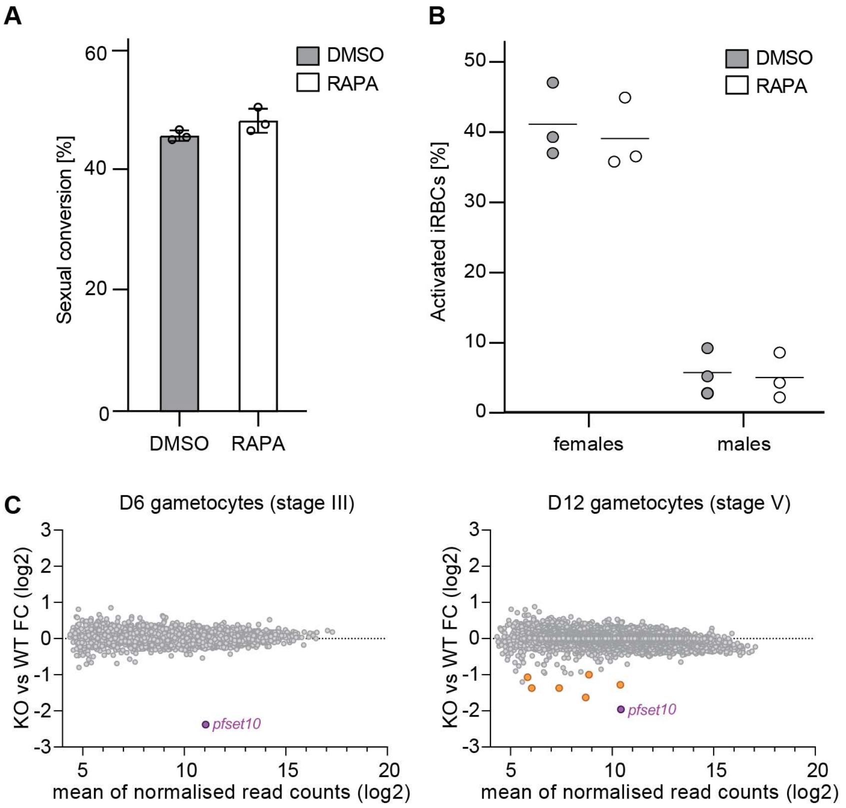


**Figure S8. PfSET10 plays no obvious role in gametocytogenesis and gamete activation. (A)** Quantification of sexual conversion rates assessed by α-Pfs16 IFAs on the progeny (30-38 hpi, stage I gametocytes) of DMSO- and RAPA-treated parasites induced for sexual commitment by mFA in the previous cycle. The means ±s.d. (error bars) of three biological replicates are shown, with circles representing individual values. At least 150 cells have been scored for each condition and replicate. (**B**) Quantification of female and male gametocyte activation rates comparing DMSO- and RAPA-treated mature stage V parasites. The percentages of activated gametocytes among all gametocytes measured in the female activation assay (Pfs25 surface expression; left) and male activation assay (exflagellation; right) are shown on the y-axis. The individual values obtained from three independent replicates are shown as circles, with the means indicated by a horizontal line. **(C)** MA plots showing mean normalized read counts (x-axis) plotted against the fold change in gene expression between control and PfSET10 KO parasites (y-axis) for all genes in day 6 (stage III; left panel) and day 12 gametocytes (stage V; right panel). Coloured circles represent differentially expressed genes identified by DESeq2 analysis (log2 fold change (FC) ≥1; adjusted p-value <0.01). Values represent the mean of three biological replicate RNA-seq experiments performed for each condition and TP. WT, DMSO-treated control gametocytes; KO, RAPA-treated PfSET10 KO gametocytes.

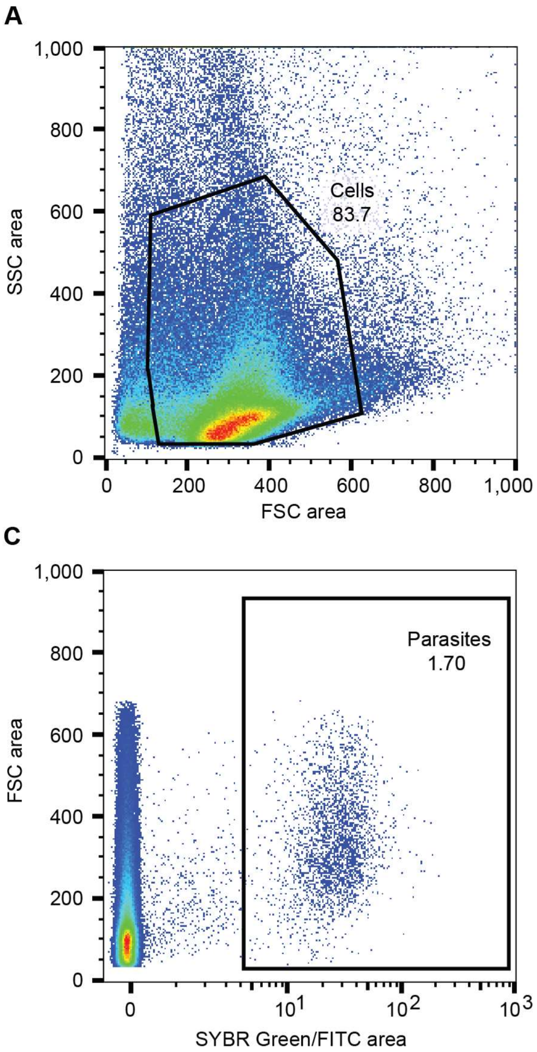


**Figure S9. Flow cytometry gating strategy. (A-C)** The gating strategy aimed at distinguishing cells from small debris (FSC-A vs. SSC-A) (A)**,** removing measurements with more than one cell per droplet (FSC-A vs. FSC-H) (B), and identifying iRBCs based on their SYBR Green I signal intensity (FSC-A vs. FITC-H) (C). Representative flow cytometry data is shown. FSC, forward scatter; SSC, side scatter; FITC, fluorescein isothiocyanate; A, area; H, height.

**Table S1: All oligonucleotides used in this study.**

| Name | sequence (5'-3') | used for | Cell line/plasmid |
|---|---|---|---|
| sgRNA_SET10_Cterm_fw | tattagacatgttttaaatagaca | cloning | pHF-gC_SET10-Cterm |
| sgRNA_SET10_Cterm_rv | aaactgtctatttaaaacatgtct | cloning | pHF-gC_SET10-Cterm |
| sgRNA_SET10_S420_5_fw | tattAATGCGAATCGAAGAAAAAG | cloning | pBF-gC_SET10-Nterm |
| sgRNA_SET10_S420_5_rv | aaacCTTTTTCTTCGATTCGCATT | cloning | pBF-gC_SET10-Nterm |
| sgRNA_v08-g4_F | tattgatgaatgaattgttagaaa | cloning | pYF_gC-v08 |
| sgRNA_v08-g4_R | aaactttctaacaattcattcatc | cloning | pYF_gC-v08 |
| NCS_F | GGTTCTGGAAAaaGACGtTGG | cloning | pMBP_NCS |
| NCS_R | GGATCCACCACTACTAGTC | cloning | pMBP_NCS |
| N0_F | GACTAGTAGTGGTGGATCCACAAGTAAAGGAGAAG | cloning | pMBP_NCS |
| N1 | TGTGTAACTCATGTGTtGCTGGaAGtGAaGCCATGTTATCTTCTTCTCCTTTACTTGTGG | cloning | pMBP_NCS |
| N2 | AGCaACACATGAGTTACACATCTTTGGaTCtATCAACGGTGTaGAtTTcGACATGGTtGG | cloning | pMBP_NCS |
| N3 | TAAtTCCTCATAACCATCATTaGGATTaCCaGTtCCtTGACCaACCATGTCgAAaTCtAC | cloning | pMBP_NCS |
| N4 | CtAATGATGGTTATGAGGAaTTAAACCTtAAaTCaACCAAGGGTGAtCTtCAGTTCTCCC | cloning | pMBP_NCS |
| N5 | TGATGGAAtCCATAaCCGATATGAGGGACtAaAATCCAaGGGGAGAACTGaAGaTCACCC | cloning | pMBP_NCS |
| N6 | ATCGGtTATGGaTTCCATCAaTACCTtCCaTAtCCTGACGGtATGTCaCCTTTCCAaGCa | cloning | pMBP_NCS |
| N7 | TTGTtCtATGaACTTGaTAaCCaGAtCCATCTACCATaGCtGCtTGGAAAGGtGACATaC | cloning | pMBP_NCS |
| N8 | GGtTAtCAAGTtCATaGaACAATGCAGTTTGAAGATGGTGCtTCaCTTACTGTaAACTAt | cloning | pMBP_NCS |
| N9 | tGCtTCTCCTTTGATGTGaCTTCCCTCaTAGGTGTAaCGaTAGTTtACAGTAAGtGAaGC | cloning | pMBP_NCS |
| N10 | GtCACATCAAAGGAGAaGCaCAaGTtAAaGGtACTGGTTTtCCTGCTGACGGTCCTGTaA | cloning | pMBP_NCS |
| N11 | TTtGAtCTaCACCAGTCaGCaGCaGTaAGtGAGTTGGTCATtACAGGACCGTCAGCAGGa | cloning | pMBP_NCS |
| N12 | GCtGACTGGTGtAGaTCaAAGAAaACTTACCCaAACGACAAAACCATtATCAGTACaTTT | cloning | pMBP_NCS |
| N13 | CGGTAtCttTTtCCATTTCCAGTGGTGTAACTCCAtTTAAAtGTACTGATaATGGTTTTG | cloning | pMBP_NCS |
| N14 | GGAAATGGaAAaaGaTACCGttcaACTGCaaGaACaACCTACACtTTTGCtAAaCCAATG | cloning | pMBP_NCS |
| N15 | GGAAaACGTACATtGGtTGGTTCTTtAaATAGTTAGCtGCCATTGGtTTaGCAAAaGTGT | cloning | pMBP_NCS |
| N16 | CCAaCCaATGTACGTtTTCCGTAAGACaGAaCTtAAaCAtTCaAAGACCGAG | cloning | pMBP_NCS |
| N17 | CtGTAAAaGCCTTTTGCCACTCCTTGAAGTTtAaCTCGGTCTTtGAaTGtTTaA | cloning | pMBP_NCS |
| N18 | GTGGCAAAAGGCtTTTACaGATGTaATGGGAATGGATGAACTATACAAAGG | cloning | pMBP_NCS |
| N19_R | CCAaCGTCttTTTCCAGAACCTGTACCTGATCCACCTTTGTATAGTTCATCCATTCC | cloning | pMBP_NCS |
| Blox_1F | GTTGGCCGATTCATTAATGGTACTTTTGTTGGTAGAG | cloning | pD_SET10BxNCS |
| Blox_1R | GTATTATCTTTCATGCTTCCACTTGTAGACATAGTTCTTTTTCTTG | cloning | pD_SET10BxNCS |
| Blox_2F | GTGGAAGCATGAAAGATAATACAGTACCattaaaattaatag | cloning | pD_SET10BxNCS |
| Blox_2R | tttacctaaaccttcttgttcaaataattc | cloning | pD_SET10BxNCS |
| Blox_3F | tttgaacaagaaggtttaggtaaataaaaaaaataatatacaATAACTTCG | cloning | pD_SET10BxNCS |
| Blox_3R | cttgataaataAggagctaaaagaatataaaatatataaatatata | cloning | pD_SET10BxNCS |
| Blox_4F | cttttagctccTtatttatcaagatggg | cloning | pD_SET10BxNCS |

| | | | |
|---|---|---|---|
| Blox_4R | CCACCAGAACCtttttcagctgatcttaatgatatttc | cloning | pD_SET10BxNCS |
| Blox_5F | ctgaaaaaGGTTCTGGTGGATCCACAAGTAAAG | cloning | pD_SET10BxNCS |
| Blox_5R | CCATATCCTCGAGATTAaggttctctttgTCCtTGTGG | cloning | pD_SET10BxNCS |
| Blox_6F | cctTAATCTCGAGGATATGGCAGCTTAATG | cloning | pD_SET10BxNCS |
| Blox_6R | catatattatttggtaacCCTGAAGAAGAAAAGTCC | cloning | pD_SET10BxNCS |
| Blox_7F | ggttataatataatatgcacaactg | cloning | pD_SET10BxNCS |
| Blox_7R | cCTCTTCGCTATTACGCCagatattattctttacatgaaaatttaatttcc | cloning | pD_SET10BxNCS |
| SET10_x_S420_5HB_fw | cgttggccgattcattaatgTTAGAACATACCAAGGATATGGG | cloning | pD_SET10xS420 |
| SET10_x_S420_5HB_rv | tCTaCgtACtTTCTTtTTcTTCTTTTCATATATAGTATCCTTAATTTTGCTTG | cloning | pD_V08-2TY-2A-BSD |
| SET10_x_S420_fw | AAgAAaAAGAAaGTacGtAGaAGgtaaataaaaaaaataatatacaATAACTTCG | cloning | pD_V08-2TY-2A-BSD |
| SET10_x_S420_rw | CCaCgcTTaCgTCtgTTaGCgTTTACTTCgAAgctaaaagaatataaaatatataaatatatatatatac | cloning | pD_V08-2TY-2A-BSD |
| SET10_x_S420_3HB_fw | CtAAcaGAcGtAAgcGtGGaGCtGGtCAAAAGGGCGTATTTTCTGATG | cloning | pD_V08-2TY-2A-BSD |
| SET10_x_S420_3HB_rv | cctcttcgctattacgccagCTTTGGTACAGCATTATCATCC | cloning | pD_V08-2TY-2A-BSD |
| v08_1F | cgttggccgattcattaatggtgattatgacatttacgatg | cloning | pD_V08-2TY-2A-BSD |
| v08_1R | tatGtcACTAataggGtattcTttCtcAaGTaattcattcatcttattattatgaagagc | cloning | pD_V08-2TY-2A-BSD |
| v08_2F | GATCTTGGTTtGTgTGaACtTCtgatccGatGttccatatGtcACTAataggGtattcTt | cloning | pD_V08-2TY-2A-BSD |
| v08_2R | GTtCAcACaAACCAAGATCCAtTaGACggtactGAGGTCCATACTAACCAAGATCCttt | cloning | pD_V08-2TY-2A-BSD |
| v08_3F | gttaataaacttcctcttccttcaccacttccaTCtaaaGGATCTTGGTTAGTATGG | cloning | pD_V08-2TY-2A-BSD |
| v08_3R | gaaggaagaggaagtttattaacatgtggagatgtagaagaaaatccaggaccaatggca | cloning | pD_V08-2TY-2A-BSD |
| v08_4F | aaatccaggaccaatggcacctttgtctcaag | cloning | pD_V08-2TY-2A-BSD |
| v08_4R | ctatatttgtattaAccctcccacacataaccag | cloning | pD_V08-2TY-2A-BSD |
| v08_5F | gtgtgggagggTtaatacaaatatagatgtatatatgtatgtatatatttttttg | cloning | pD_V08-2TY-2A-BSD |
| v08_5R | cctcttcgctattacgccagcataacatattcatagtatatagc | cloning | pD_V08-2TY-2A-BSD |
| CS0_F | GGTGGATCCGGTACAGGTTCTGGAAAaaGACGtTGGAAAAAGAATTTCATcGC | cloning | pCS |
| CS1 | CactGGATGAaATTTTCTTAAAaCgGTTtGCaGCactaACaGCgATGAAATTCTTTTTCC | cloning | pCS |
| CS2 | GtTTTAAGAAAATtTCATCCagtGGtGCAttaGGTGGAAGTGGTAGTGGAGGAggtatgg | cloning | pCS |
| CS3 | CtTCaACtACGTGtCCACCtctccatccggttgtcttttcatccataccTCCTCCACTAC | cloning | pCS |
| CS4 | GTGGaCACGTaGTtGAaGGttTaGCaGGaGAGtTaGAaCAatTaaGaGCaAGatTaGAaC | cloning | pCS |
| CS5_R | GGTGCTCGAGTTAtggttctctttgTCCtTGtGGGTGaTGtTCtAatCTtGCtCttAatTG | cloning | pCS |
| pYF_F1 | GCTTGTCGACGGAGCTC | cloning | pYF_gC |
| pYF_R1 | GGCTGTCATTGATATATTTCTATTAGGTATTtattattataaaatataaatc | cloning | pYF_gC |
| pYF_F2 | CCTAATAGAAATATATCAATGACAGCCAGTTTAACTACC | cloning | pYF_gC |
| pYF_R2 | GATCCTGAaccAATGCTGTTCAACTTCCC | cloning | pYF_gC |
| pYF_F3 | CAGCATTggtTCAGGATCaGTGACAG | cloning | pYF_gC |
| pYF_R3 | gtagccatgatagcactacc | cloning | pYF_gC |
| SET C out F | cttataggagtagaattaggaaagacg | PCR on gDNA | 3D7 & NF54/DiCre/SET10-BirA*xNCS |
| SET C in F | ggatatggcagcttaatgttcg | PCR on gDNA | 3D7 & NF54/DiCre/SET10-BirA*xNCS |
| SET C in R | cgaacattaagctgccatatcc | PCR on gDNA | 3D7 & NF54/DiCre/SET10-BirA*xNCS |
| SET C out R | ccttaacatacagtgaactttaataatatatgag | PCR on gDNA | 3D7 & NF54/DiCre/SET10-BirA*xNCS |
| SET C WT R | cttatatttcaagccagttgtaatttg | PCR on gDNA | 3D7 & NF54/DiCre/SET10-BirA*xNCS |
| SETx out F | ATGAATAAAGAAAGAATGGATGACG | PCR on gDNA | 3D7 & NF54/DiCre/SET10-mNG-iKO |

| | | | |
|---|---|---|---|
| SETx in F | GAAGTAAAcGCtAAcaGAcG | PCR on gDNA | 3D7 & NF54/DiCre/SET10-mNG-iKO |
| SETx in R | gtctgTTaGCgTTTACTTCg | PCR on gDNA | 3D7 & NF54/DiCre/SET10-mNG-iKO |
| SETx out R | GGTTATTACTCAAATTAGGATCTTCC | PCR on gDNA | 3D7 & NF54/DiCre/SET10-mNG-iKO |
| SETx WT R | attcgcatttacttcaaaactcc | PCR on gDNA | 3D7 & NF54/DiCre/SET10-mNG-iKO |
| v08 out F | cagtggtattgatccaacaagtgc | PCR on gDNA | 3D7 & NF54/DiCre/SET10-mNG-iKO/V08-2xTY-BSD |
| V08 in F | ctctggttatgtgtgggagg | PCR on gDNA | 3D7 & NF54/DiCre/SET10-mNG-iKO/V08-2xTY-BSD |
| v08 in R | cttcctcttccttcaccacttcc | PCR on gDNA | 3D7 & NF54/DiCre/SET10-mNG-iKO/V08-2xTY-BSD |
| v08 our R | gatggtacaaacaatataattaaaataatctcatatggtag | PCR on gDNA | 3D7 & NF54/DiCre/SET10-mNG-iKO/V08-2xTY-BSD |
| v08 WT R | ctatatttgtattatatattccatatatctg | PCR on gDNA | 3D7 & NF54/DiCre/SET10-mNG-iKO/V08-2xTY-BSD |
| Bb R | ATTCGCCATTCAGGCTGC | PCR on gDNA | 3D7 & NF54/DiCre/SET10-mNG-iKO/V08-2xTY-BSD |
| gapdh int F | aatggcagtaacaaaacttgg | PCR on gDNA | positive control |
| gapdh int R | cttagttgttagtaatgtgtacgg | PCR on gDNA | positive control |

**Dataset S1: Quantification of *var* gene transcripts in 3D7/DiCre/SET10-mNG-iKO/V08-2xTYBSD ring stage parasites (10-16 hpi) by RNA-seq.** Columns A-B: Gene ID and gene description (www.plasmodb.org; release ver. 66). Columns C-Q: Normalised read counts quantified in each individual replicate samples. R-WT-G0, control parasites selected on BSD-S-HCl for *v08-2xty* expression; R-WT-G2/R-WT-G10, DMSO-treated control parasites harvested two/ten generations after release from BSD-S-HCl selection pressure, respectively; R-KO-G2/R-KO-G10, RAPA-treated PfSET10 KO parasites harvested two/ten generations after release from BSD-S-HCl selection pressure, respectively. Columns R-AA: Mean normalized read counts and s.d. for each of the five triplicate samples. Columns AB-AE: mean log2 fold change (FC) in *var* transcript abundance between the R-WT-G2/R-WT-G10 and R-KO-G2/R-KO-G10 samples compared to the R-WT-G0 control reference sample.

**Dataset S2: Differential gene expression by DESeq2 analysis between 3D7/DiCre/SET10-mNG-iKO/V08-2xTY$^{BSD}$ DMSO control and RAPA-treated PfSET10 KO parasites at four TPs during the IDC.** Columns A-B: Gene ID and gene description (www.plasmodb.org; release ver. 66). Columns C-Y: Normalised read counts quantified in each individual replicate sample. R/T/ES/LS-WT-G2, DMSO-treated control parasites harvested two generations after release from BSD-S-HCl selection pressure from synchronous ring stages (R, 10-16 hpi), trophozoites (T, 22-28 hpi), early schizonts (ES, 30-36 hpi) and late schizonts (ES, 38-44 hpi), respectively; R/T/ES/LS-KO-G2, RAPA-treated PfSET10 KO parasites harvested two generations after release from BSD-S-HCl selection pressure from synchronous ring stages (R, 10-16 hpi), trophozoites (T, 22-28 hpi), early schizonts (ES, 30-36 hpi) and late schizonts (LS, 38-44 hpi), respectively. Columns Z-AG: Mean normalized read counts for each of the eight triplicate samples. Columns AH-AW: mean normalized read counts (log2) of all six R (red), T (green), ES (AP) and LS (purple) samples; mean log2 fold change (FC) in transcript abundance between RAPA-treated PfSET10 KO and DMSO control R, T, ES and LS samples; Benjamini-Hochberg adjusted p-value; genes significantly upregulated (up; log2 FC ≥1) or downregulated (down; log2 FC ≤-1) (adjusted p-value <0.01) in R, T, ES and LS samples. Columns AX-AY: TP of minimal and maximal expression. 1, R; 2, T; 3, ES; 4, LS. Columns AZ-BC: genes with mean normalized read counts ≤20 in either one or both R, T, ES or LS triplicate samples.

**Dataset S3: Differential gene expression by DESeq2 analysis between 3D7/DiCre/SET10-mNG-iKO/V08-2xTY$^{BSD}$ DMSO control and RAPA-treated PfSET10 KO gametocytes.** Columns A-B: Gene ID and gene description (www.plasmodb.org; release ver. 66). Columns C-N: Normalised read counts quantified in each individual replicate sample. D6/D12-WT, DMSO-treated control stage III gametocytes (D6, day 6) and stage V gametocytes (D12, day 12), respectively; D6/D12-KO, RAPA-treated PfSEt10 KO stage III gametocytes (D6, day 6) and stage V gametocytes (D12, day 12), respectively. Columns O-R: Mean normalized read counts for each of the four triplicate samples. Columns S-Z: mean normalized read counts (log2) of all six D6 (red) and D12 (green) samples; mean log2 fold change (FC) in transcript abundance between RAPA-treated PfSET10 KO and DMSO control D6 and D12 samples; Benjamini-Hochberg adjusted p-value; genes significantly upregulated (up; log2 FC ≥1) or downregulated (down; log2 FC ≤-1) (adjusted p-value <0.01) in D6 and D12 samples. Columns AA-AB: genes with mean normalized read counts ≤20 in either one or both D6 or D12 triplicate samples.
